# Revealing and evaluation of antivirals targeting multiple druggable sites of RdRp complex in SARS-CoV-2

**DOI:** 10.1101/2023.07.24.550324

**Authors:** Ruchi Rani, Sanketkumar Nehul, Shweta Choudhary, Anushka Upadhyay, Gaurav Kumar Sharma, Pravindra Kumar, Shailly Tomar

## Abstract

SARS-CoV-2 RNA-dependent RNA polymerase (RdRp) complex consisting of nsp12, nsp7, and nsp8 as the key enzyme for viral genome replication and is a proven antiviral drug target. In this study, molecular interactions of nsp7 and nsp8 with nsp12 and the active site of nsp12 were coterminously targeted using *in-silico* screening of small molecule libraries to identify potential antivirals. Surface plasmon resonance (SPR) based assay using purified nsp7 and nsp8 proteins was developed, and the binding of identified molecules to targets was validated. The antiviral efficacy of identified small molecules was evaluated using cell-based assays, and potent antiviral effect with EC_50_ values of 0.56 μM, 0.73 μM, and 2.8 μM was demonstrated by fangchinoline, cepharanthine, and sennoside B, respectively. Further *in vivo,* investigation using *hACE2* mice is being conducted. This is the first study that targets multiple sites in the RdRp complex of SARS-CoV-2 using a structure-based molecular repurposing approach and suggests potential therapeutic options for emerging variants of SARS-CoV-2.

## Introduction

SARS-CoV-2, a novel β-coronavirus (CoV), was initially identified in December 2019 and quickly emerged as the causative agent behind the global COVID-19 pandemic. Despite its relatively low mortality rate, SARS-CoV-2 remains a significant research focus due to its remarkable transmissibility and widespread prevalence (Chan et al., 2020). The primary cause of COVID-19’s spread was lack of effective licensed antiviral drug molecules and effective vaccines, which dramatically increased infections. As of now, vaccination campaigns are gaining momentum worldwide; there are encouraging signs of relief even in regions heavily impacted by the pandemic. Nevertheless, the emergence of new vaccine-resistant strains of CoV remains a concern, with the constant evolution of the virus (Lopez Bernal et al., 2021; Wilson et al., 2020). Furthermore, there is the possibility that certain vaccinated individuals may not generate a robust immune response against new strains of SARS-CoV-2 upon subsequent exposure, leading to insufficient production of neutralizing antibodies. With a better understanding of the viral genomic structure, scientific research studies are concentrating on repurposing therapeutic drugs that target the virus conserved proteins, which are essential for the virus life cycle or for the viral replication machinery.

Coronaviruses utilize a complex of non-structural proteins (nsps) known as the RdRp complex to facilitate the replication and transcription processes of their RNA genome within host cells (Wu et al., 2022). This RdRp complex comprises three distinct nsp subunits: nsp12 (catalytic subunit), and the two accessory subunits: nsp7 and nsp8 (Gao et al., 2020; Hillen et al., 2020). The catalytic subunit nsp12, exhibits limited activity, and its proper functioning relies on the presence of nsp8 and nsp7 proteins (Te Velthuis et al., 2012; Q. Wang et al., 2020). While the specific role of SARS-CoV-2 nsp7 is unknown, it is observed that the two copies of nsp8 make elongated helices that act as critical interfaces for RNA binding (Hillen et al., 2020; Te Velthuis et al., 2012). These elongated helices may also act as a stabilizing component for the RNA template/product duplex generated during replication within the catalytic core of nsp12 (Hillen et al., 2020; Wang et al., 2020). Additionally, nsp8 self-assembly into hexa-dodecameric complexes lead to polymerase/primase activity (Te Velthuis et al., 2012). This suggests that nsp7-nsp8 may have two distinct and structurally independent functions but also play crucial roles in enhancing RdRp template binding and processivity with catalytic core, nsp12 (Kirchdoerfer and Ward, 2019). Various crystal structures of the RdRp of SARS-CoV-2 have been successfully obtained, revealing its conformation both in the RdRp complex state (Kirchdoerfer and Ward, 2019) and in complex with RNA template-product duplexes (Yan et al., 2020). These crystal structures provide valuable insights into the binding interactions and conformational changes that occur within the RdRp complex, highlighting its role in facilitating viral genome replication and transcription.

Remdesivir (Veklury^®^), a nucleoside analog, stands as the sole FDA-approved drug against SARS-CoV-2 RdRp for treating patients with COVID-19 (Paladugu and Donato, 2020; Yin et al., 2020). By inhibiting the RdRp of CoVs, remdesivir has demonstrated antiviral activity in both cell-based and *in vivo* studies (Wang et al., 2020). Nguyenla et al. reported a notable antiviral efficacy of remdesivir, demonstrating an EC_50_ of 3 ± 0.6 μM (Nguyenla et al., 2022), which aligns with various reported values ranging from 0.6 to 11 μM in Vero-E6 cells (Pruijssers et al., 2020; Wang et al., 2020). While robust effects of remdesivir has been observed in non-human primate models of COVID-19 (Dobrovolny, 2020), but exhibits limited efficacy in humans (Goldman et al., 2020; Olalla, 2020; Spinner et al., 2020). The complex pharmacokinetic profile of remdesivir, coupled with its relatively low potency, may restrict its efficacy due to diffusion-driven distribution to the target tissue (Jorgensen et al., 2020) (Sun, 2020). Therefore, there is an urgent need of alternative repurposed drugs to ameliorate treatment of SARS-CoV-2 infections. Additionally, other compounds such as favipiravir (Du and Chen, 2020; Shannon et al., 2020) and molnupiravir (Agostini et al., 2019; Sheahan et al., 2020) are also currently under investigation for their potential to disrupt viral replication, aiming to enhance antiviral effectiveness.

Our study aims to assess potential drugs by targeting catalytic site and the protein-protein interface residues (PPI) in the RdRp complex of SARS-CoV-2, using biophysical and cell culture-based analyses. Among the compounds selected, fangchinoline, cepharanthine and sennoside B have emerged as highly promising candidates, demonstrating EC_50_ values of 0.56 ± 0.14 μM, 0.73 ± 0.3 μM and 2.80 ± 0.61 μM, respectively in Vero cells. These values indicate their superior antiviral potency compared to the FDA-approved drug remdesivir. Consequently, these compounds hold significant potential for further evaluation in *in vivo* and clinical trials against SARS-CoV-2. As their promising potency suggests, the compounds are currently undergoing further assessment through *in vivo* studies.

## Methodology

### Software, Libraries, and Database

The PyRx 0.8 (O’Boyle et al., 2011; Dallakyan and Olson, 2015), AutoDock MGL tools 1.5.6, AutoDock Vina (Steffen et al., 2010), and LIGPLOT+ along with PyMOL 2.3.4, (Wallace et al., 1995), were utilized in this study. The Research Collaboratory for Structural Bioinformatics (RCSB) Protein Data Bank (PDB) database (Berman et al., 2000) served as the primary source of protein structural information. For screening purposes, three drug libraries were selected: Library of Pharmacologically Active Compound (LOPAC^1280^) from Sigma, the Food and Drug Administration (FDA) approved library and natural product library (NPL) from Selleckchem, and virtual screening was performed on a macOS Mojave workstation.

### Preparation of Ligands and Proteins

The three-dimensional structures of RdRp protein complexes of SARS-CoV-2 (nsp12, nsp7 and nsp8: PDB ID 6M71) were retrieved from RCSB PDB in .pdb format. Three drug libraries—FDA-approved drug library, NPL from Selleckchem (2747 and 2370 compounds, respectively), and LOPAC1280 from Sigma (1280 compounds)—were retrieved in.sdf format in order to identify potential antiviral compounds. The virtual screening procedure for proteins and ligands was followed as mentioned in Rani et al., 2022.

### Computer-aided structure-based virtual screening

To investigate the binding affinities of the compounds, three different sites were targeted in the nsp12 protein in two steps. In the first step of screening, all the molecules from the different libraries were screened against the catalytic active site of RdRp of SARS-CoV-2 protein using PyRx 0.8 accompanied by AutoDock Vina. The grid centre points were set at X = 25, Y = 21, Z = 25, and box dimensions were set as 119 Å ×117 Å × 128 Å with exhaustiveness 8.

Following the initial screening, the selected molecules underwent a second round of virtual screening targeting the interactive residues of nsp12 with nsp8 and nsp7 using PyRx 0.8 accompanied by AutoDock Vina. For this purpose, the grid centre points were set at X = 47, Y = 33, Z = 34, and box dimensions were set as 116 Å × 144 Å × 138 Å, for nsp12 interacting residues for nsp8, with an exhaustiveness 8. While, the grid centre points for nsp12 interacting residues for nsp7 were set at X = 39, Y = 33, Z = 23, and box dimensions were set as 112 Å × 102 Å × 149 Å with exhaustiveness 8 (**Supplementary table 1**).

### Molecular Docking

Molecular docking studies were performed for a set of nucleotide monophosphates (NMPs) including AMP, GMP, CMP, and UMP, as well as NTPs such as ATP, GTP, CTP, and UTP. Additionally, positive control compounds like remdesivir, sofosbuvir, ribavirin, galidesivir, and favipiravir were included in the docking analysis. The binding affinity of these positive controls were compared with the virtually screened compounds, and molecules with binding energy (B.E.) ≥9 kcal/mol against the catalytic site residues of nsp12 were selected for further screening. Additionally, the selected compounds were subjected to screening molecules with B.E. ≥8 kcal/mol against the interface residues of nsp12. We have also screened all the compounds against the nsp8 and nsp7 interface residues with nsp12. For detailed analysis, AutoDock Vina algorithm was employed for molecular docking of the selected compounds (Trott and Olson, 2009). The grid parameters set for molecular docking studies are listed in **Supplementary table 1**.

Two-dimensional representation of interactions between SARS-CoV-2 RdRp proteins and ligands were prepared using PyMOL 2.3.4 and LIGPLOT^+^.

### Preparation of compounds stock solutions

Amentoflavone (Catologue No. 21779), chikusetsusaponin iva (Catologue No. 34832), xifaxan (Catologue No. 16131), natacyn (Catologue No. 11634), sennoside B (Catologue No. 2841), rapamycin (Catologue No. 13346), dihydroergotamine methanesulfonate (Catologue No. 23847), cepharanthine (Catologue No. 19648), fangchinoline (Catologue No. 29243), and doramectin (Catologue No. 19467) were procured from Cayman, USA. Stock solutions of all the compounds were prepared using dimethyl sulfoxide (DMSO) obtained from Sigma-Aldrich. Prior to the experiments, further working dilutions of the compounds were prepared in culture media for cell culture experiments and in buffers for biophysical assays.

### Purification of SARS-CoV-2 nsp7 and nsp8 cofactors

The full-length nsp8 (encoding 1-198 residues) and nsp7 (encoding 1-84 residues) were received from Krogan’s Lab which were cloned into pET28c(6X-his-nsp8) and pET28c(6X-his-nsp7) for the recombinant protein expression of nsp8 and nsp7 in *E. coli* cells.

Positive clones of pET28c-(6X-his-nsp8) and pET28c-(6X-his-nsp7) were selected for the expression of recombinant nsp8 and nsp7 protein by transforming chemically competent *E. coli* BL21 (DE3) cells. The cells were spread onto the Luria-Bertani (L.B.) agar plate containing kanamycin (50 μg/mL). The plates were incubated at 37 °C overnight. A single colony was inoculated into kanamycin positive (50 μg/mL) L.B. broth and grown at 37 °C overnight.

Bulk expression of recombinant proteins was performed individually by inoculating 10 mL of the primary culture of transformed BL21 cells in 1 L of L.B. and incubated at 37 °C in an incubator shaker for 3 h until optical density (O.D_600_) reached 0.6. The culture was induced with 0.1 mM isopropyl-β-D-1-thiogalactopyranoside (IPTG) followed by an incubation period of 20 h at 18 °C with continuous shaking at 180 rpm. The culture was then harvested at 6000 rpm at 4 °C for 10 min. The pellets were resuspended in lysis buffer containing 50 mM Tris (pH 8), 200 mM NaCl, and 1 mM phenylmethylsulfonyl fluoride (PMSF). Cells were lysed on ice using a sonicator. The cell lysate was centrifuged at 13,000 rpm for 90 min at 4 °C. The supernatant was loaded onto Ni^+2^-NTA column beads and unwanted proteins (impurities) were eliminated by subsequent washing with a binding buffer containing 200 mM NaCl and 20-40 mM imidazole. A gradient of 50-500 mM imidazole was performed to elute proteins. Fractions containing purified proteins were pooled and dialyzed against 50 mM HEPES (pH 8.0) buffer containing 150 mM NaCl, for imidazole removal. The pooled elution fractions of both the proteins was analysed on 12% sodium dodecyl sulfate (SDS)-polyacrylamide gel electrophoresis (PAGE). The purified fractions of the proteins were concentrated up to 3-4 mg/mL in 30-kDa molecular weight cut-off Amicon concentrator (Millipore, USA) for further experiments.

### Surface Plasmon Resonance (SPR) based binding kinetics

The binding kinetic studies of selected compounds to nsp8 or nsp7 proteins was performed by SPR using a Biacore T200 system (Biacore Inc., Uppsala, Sweden). Before starting the experiment, the Biacore T200 instrument was washed and cleaned using the manufacturer’s standard desorb and sanitize procedure protocol to get good-quality SPR sensorgram data. For optimization, the instrument was primed four-five times after the procedure was finished and left overnight in standby mode. The temperature of the sample compartment for binding and kinetic of all identified compounds experiments were maintained at 25 °C. The Ni-NTA sensor chip was pre-conditioned with degassed and filtered 50 mM HEPES (pH 8.0), and 150 mM NaCl buffer prior to His-tag nsp8/nsp7 immobilization while every flow channel has its own reference channel.

The purified His-tagged nsp8 or nsp7 protein was diluted to 20 μg in degassed and filtered 50 mM HEPES (pH 8.0), and 150 mM NaCl buffer for immobilization on Ni-NTA sensor chips. The selected compounds solutions with different increasing concentrations were flowed over the surface of active (immobilized his-tagged protein) and reference (blank) channels at a flow rate of 30 μL/min with a dissociation time of 90 s and contact time of 60 s. The dissociation phase was followed by regeneration of surface with 350 mM EDTA.

SPR data were obtained using the Biacore T200 control software and analyzed with the Biacore T200 evaluation software (Software version 3.2). The obtained response unit (R.U.) signal was determined by subtracting the R.U. of the reference cell from the R.U. of the active flow cell (immobilised His-tagged nsp8 or nsp7 proteins). The values of the association constant (K_on_), dissociation constant (K_off_), and affinity (KD) for all the selected compounds were calculated by fitting the data to 1:1 Langmuir binding model.

### Cells and SARS-CoV-2 virus

Vero cell line was obtained from NCCS, Pune and the cells were subsequently maintained in Dulbecco’s modified eagle medium (DMEM, Gibco) augmented with 10% heat-inactivated fetal bovine serum (FBS, Gibco), PenStrep antibiotic (50 U mL^−1^ penicillin, and 50 μg mL^−1^ streptomycin). The cells were cultured at 37 °C and 5% CO_2_ in a humidified incubator.

The SARS-CoV-2/Human/IND/CAD1339/2020 strain was isolated from COVID-19 positive patient and subsequently inoculated on Vero cells and confirmed via sequencing as described by Dhaka et al., 2022. The aliquots of confirmed SARS-CoV-2 virus were prepared and stored at -80 °C for research purpose. Virus titer was estimated using the median tissue culture infectious dose (TCID_50_) assay.

### Cell viability assay

The cytotoxic effects of the identified compounds on Vero cells were determined using the standard MTT (3-(4, 5-dimethyl thiazol-2yl)-2, 5-diphenyl tetrazolium bromide) assay. Briefly, in a 96-well plate, 1×10^5^ cells were seeded in 100 µL of medium per well. Various concentrations of compounds ranging 1.25 µM to 200 µM were added onto Vero cells at 90% confluence and incubated for 48 h at 37 °C. Each treatment was replicated three times. After an incubation of 48 h, the medium was removed from all the wells, followed by the addition of 20 µL of MTT solution (5 mg/mL) to each well. The plate was incubated for 4 h in dark in incubator at 37 °C and 5% CO_2_. Later, 110 µl of DMSO was added per well to dissolve the formazan crystals. The cell viability was determined by quantifying the resulting absorbance at 570 nm using a multi-mode microplate reader (BioTek Instruments, Inc.), with cell control (CC) serving as a blank. The absorbance of formazan produced in control cells was considered as 100% viability. The data is presented as a mean percentage of viable cells compared to their respective controls.

### Assessment of Antiviral Activity

On a day prior to infection for antiviral assessment of compounds, Vero cells were seeded onto 24-well plate and then incubated at 37 °C and 5% CO_2_. The next day, the media was removed and the cells were pre-treated with various concentrations of compounds for 3 h. Wash with 1X phosphate buffer saline (PBS) was given and subsequently the cell monolayer was infected with SARS-CoV-2/Human/IND/CAD1339/2020 virus at 0.01 MOI, with gentle shaking every 15 min for 2 h. After discarding the inoculum, the cells were washed thrice with 1X PBS. Following infection, the compounds were added to the post-infection media, which contained DMEM with 2% FBS, and the cells were further incubated for 48 h at 37 °C and 5% CO_2_.

At 48 h post-infection (hpi), the plates were freeze-thawed and total RNA was extracted from the lysate using a HiPurA™ Viral RNA Purification Kit (HiMedia), as per manufacturer’s protocol. Viral RNA was quantified by one-step qRT-PCR using the COVISure-COVID-19 Real-Time PCR kit (Genetix) according to the manufacturer’s protocol, using the following thermocycling condition: 50 °C for 15 min, 95 °C for 2 min, 95 °C for 30 s, and 60 °C for 30 s, for 40 cycles. qRT-PCR was conducted using the Applied Biosystems 7500 Fast Real-Time PCR System, and the cycle threshold (Ct) value of the SARS-CoV-2 target gene was ascertained. Graphical representations for percentage inhibition versus concentration graph was plotted using GraphPad Prism to calculate the EC_50_ values. All experiments were performed in duplicates.

### Viral Titer by TCID_50_ assay

Antiviral potential of the selected compounds was further validated by TCID_50_ assay, and the virus titers were estimated by the established Reed and Muench method (Reed and Muench, 1938). A 96-well plate was seeded with Vero cells and the plate was incubated overnight in a humidified environment at 37 °C with 5 % CO_2_. The freeze-thawed cell lysate of infected and compound treated cells were collected at 48 hpi of the antiviral assessment was subjected to ten-fold serial dilutions. The Vero cells were incubated for 3-4 days at 37 °C and 5 % CO_2_, with daily monitoring for the appearance of a CPE. In parallel, mock treated and infected Vero cell were used as a vehicle control and samples containing only culture media were used as a cell control. Post incubation, the viral inoculum was removed, and 10% formaldehyde was used as a fixative to fix the cells at room temperature for 2 h. This was followed by staining the wells for 1 h with a solution of 0.1% crystal violet. Percentages of infected dilutions immediately above and below the 50% threshold were determined, and then TCID_50_ was calculated.

### Biosafety

All experiments involving infectious viral materials were carried out in a biosafety level-3 laboratory, at the Indian Veterinary Research Institute (IVRI), Bareilly, India, in strict compliance with all necessary permissions required for studying SARS-CoV-2.

## Results

### Identification of antivirals targeting multiple sites in the RdRp complex

The CoV replication machinery relies on many protein-protein interactions, while nsp12, nsp7, and nsp8 is the main important that plays crucial roles in the formation and functionality of the RNA synthesis machinery (Kirchdoerfer and Ward, 2019). The nsp12-nsp7-nsp8 complex is the minimum essential complex necessary for effective nucleotide polymerization, even though other viral nsps cooperate in replication and transcription machinery (Lehmann et al., 2015; Sevajol et al., 2014). However, nsp12 exhibits minimal activity on its own; the presence of its co-factors, nsp7 and nsp8, significantly enhance polymerase activity (Hillen et al., 2020; Peng et al., 2020; Subissi et al., 2014; Te Velthuis et al., 2012). Firstly, a comprehensive sequence analysis of the SARS RdRp protein and its co-factors with the SARS-CoV-2 was performed using multiple sequence alignment tools. The alignment strongly indicates the conserved nature of the RdRp complex in SARS & SARS-CoV-2 protein (Supplementary figure 1). Additionally, Wang et al. showed the binding locations of remdesivir, an RdRp-targeting drug, and the RNA template around the catalytic site residues (Wang et al., 2020) (**Figure 1a**). This compelling evidence strongly indicates that the catalytic site residues represent a viable and drugable target for intervention. Moreover, PPI interfaces are less conserved than active sites, but intriguingly, as shown in figure1a, the RdRp complex interface residues are highly conserved between SARS and SARS-CoV-2. This conservation suggests an opportunity for selective obstruction of the PPI and the conserved catalytic site of the RdRp complex.

**Figure 1:**
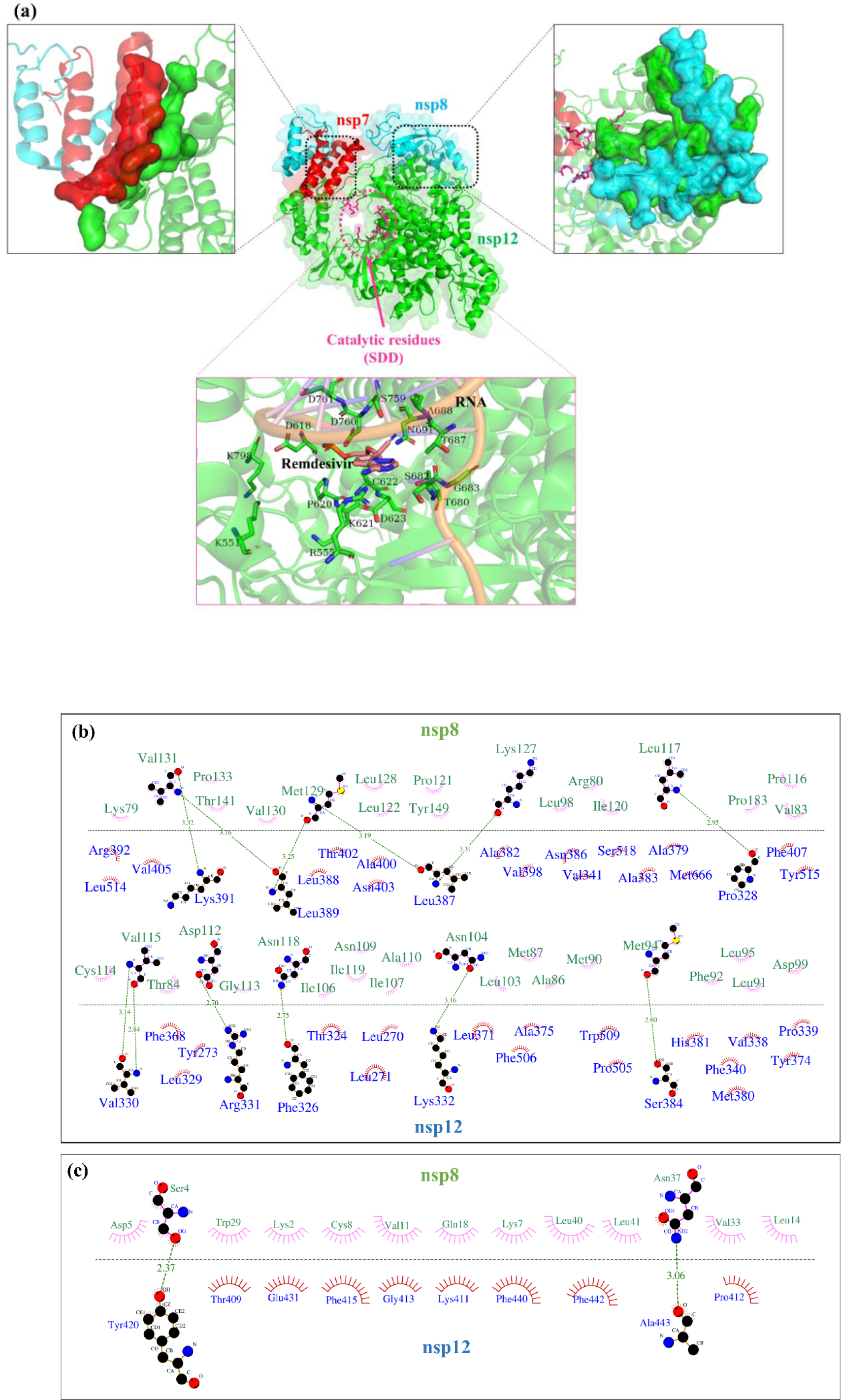
Schematic presentation of RdRp complex catalytic and interface residues. (a) Cartoon view of the nsp12-nsp8-nsp7 complexes of SARS-CoV-2 (PDB ID: 6M71) showing catalytic residues and interacting interface residues of nsp12-nsp8 and nsp12-nsp7 proteins. The nsp12 is represented in green, nsp7 in red, and nsp8 in cyan color. Interface residues between two proteins are represented in surface view with defined color. Interactive residues are mentioned in Supplementary *table 1*. Additionally, the cartoon representation highlights the interaction between nsp12 catalytic site residues (green) and the drug remdesivir (blue) and RNA template (orange) (PDB ID: 7BV2) using PyMOL. Representation of interface residues between (b) nsp8 and nsp12, and (b) nsp7 and nsp12 residues using the LIGPLOT+ analysis tool. In LIGPLOT^+^, H-bonds are represented by green dotted lines (---) with a distance in Å and hydrophobic interactions are indicated with brown curvature (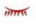) around the macromolecule residues. The interface between two proteins is depicted in a black dotted line. nsp8 protein residues are green while nsp12 residues are represented in blue color.

In this study, we focus on targeting the catalytic site and the interface residues of the RdRp complex with a same compound. The aim is to disrupt the RdRp complex’s interaction that could inhibit viral replication, ultimately gaining control over SARS-CoV-2 infection. Therefore, structure-based virtual screening of the compounds from the NPL, LOPAC library, and the FDA-approved drug library was carried out to identify cost-effective antivirals against the catalytic site and the interactive residues of SARS-CoV-2 RdRp complex. In the first round of virtual screening, the top ninety compounds from three libraries were selected based on binding affinity greater/or equal to 9 kcal/mol and greater than the positive control (**Supplementary table 2**). Nucleotide triphosphate (NTPs) (GTP, UTP, CTP, and ATP), NMPs (GMP, UMP, CMP, and AMP), and six drugs approved against different viruses of RdRp (Sofosbuvir, Ribavirin, Galidesivir, Remdesivir, Favipiravir, and Tenofovir) were used as a positive control (**Supplementary table 2**). SARS-CoV-2 RdRp incorporates these nucleotide analogues into the RNA. Once incorporated, they exert their antiviral effects through different mechanisms. Some of these analogues, such as sofosbuvir, remdesivir, and tenofovir hinder further RNA polymerisation by blocking the RdRp protein’s activity. On the other hand, favipiravir induces errors in the replicated RNA, leading to a reduction in viral replication.

The selected molecules were then sorted in increasing order of their binding affinity, and the selected compounds were subjected to another round of virtual screening against the nsp12 interface residues with nsp8 and nsp7 (**Supplementary table 3**). All the interface residues between nsp8-nsp12 and nsp7-nsp12 are represented in **figure 1b and 1c,** respectively. These interface residues were targeted in the second round of screening to select compounds whose binding affinity is ≥ 8 kcal/mol against SARS-CoV-2 nsp12 interface residues, listed in **Supplementary table 3**.

### Analysis of Molecular interactions with inhibitors

For the twenty-three ligands selected, the observed binding affinity was in the range of -8 kcal/mol to -11 kcal/mol against the catalytic and interface residues of nsp12. Furthermore, molecular docking was used to identify the best candidates from all the selected screened compounds based on the docking scores. Using the ligand availability and binding affinity of the selected compounds, we selected fourteen compounds for molecular docking to validate the procedure. The binding energy in the docking studies of all ligands against the catalytic and interface residues of nsp12, nsp8, and nsp7 are presented in **Table 1**.

**Table 1:** Binding energies (kcal/mol) of selected molecules against the catalytic site and nsp12 interactive site residues of SARS-CoV-2 proteins using AutoDock vina algorithm.

| S.No. | Ligands | Catalytic<br>site of<br>nsp12 | nsp12 interface<br>residues against |  | Targeting<br>interface residues |  |
| --- | --- | --- | --- | --- | --- | --- |
|  |  |  | nsp8 | nsp7 | nsp8 | nsp7 |
| FDA Library from Selleckchem |  |  |  |  |  |  |
| 1. | Sennoside B | -10.4 | -10.4 | -8.4 | -7.3 | -7.2 |
| 2. | Doramectin | -10.1 | -8.6 | -8.9 | -8.3 | -7.5 |
| 3. | Simeprevir | -9.3 | -9.1 | -8.7 | -7.2 | -7.3 |
| 4. | Xifaxan | -10.6 | -8.0 | -8.8 | -7.4 | -7.6 |
| Natural Product Library Compounds from Selleckchem |  |  |  |  |  |  |
| 5. | Amentoflavone | -9.5 | -8.6 | -9.1 | -7.5 | -7.8 |
| 6. | Natacyn | -9.5 | -8.5 | -8.3 | -7.1 | -7.0 |
| 7. | Chikusetsusaponin<br>Iva | -9.5 | -8.5 | -8.1 | -7.8 | -7.3 |
| 8. | Rapamycin | -9.6 | -9.0 | -9.2 | -7.5 | -7.4 |
| 9. | Cepharanthine | -9.9 | -9.8 | -9.4 | -8.0 | -7.2 |
| 10. | Fangchinoline | -9.2 | -8.0 | -9.0 | -7.2 | -7.7 |
| LOPAC from Sigma |  |  |  |  |  |  |
| 11. | Dihydroergotamine<br>methanesulfonate | -10.4 | -9.7 | -8.3 | -8.4 | -7.0 |
| 12. | KT203 | -9.4 | -8.8 | -9.2 | -7.6 | -7.8 |
| 13. | KT185 | -10.1 | -9.4 | -9.7 | -8.0 | -8.1 |
| 14. | WIN62577 | -9.7 | -9.3 | -8.7 | -7.5 | -8.1 |

Out of all, four compounds (Simeprevir, KT203, KT185, and WIN62577) have earlier been identified to target conserved pockets in other viral proteins such as Mpro proteins (Rani et al., 2020) and are giving the antiviral activity in *in vitro* study at micromolar range against SARS-CoV-2 (SARSCoV-2/Human/IND/CAD1339/2020 strain) (To be published). Therefore, interactions analysis of remaining ten compounds against nsp12, nsp8 and nsp7 protein of SARS-CoV-2 via LIGPLOT^+^ analysis tool was done (**Figure 2-4 and Table 2-3**).

**Figure 2:**
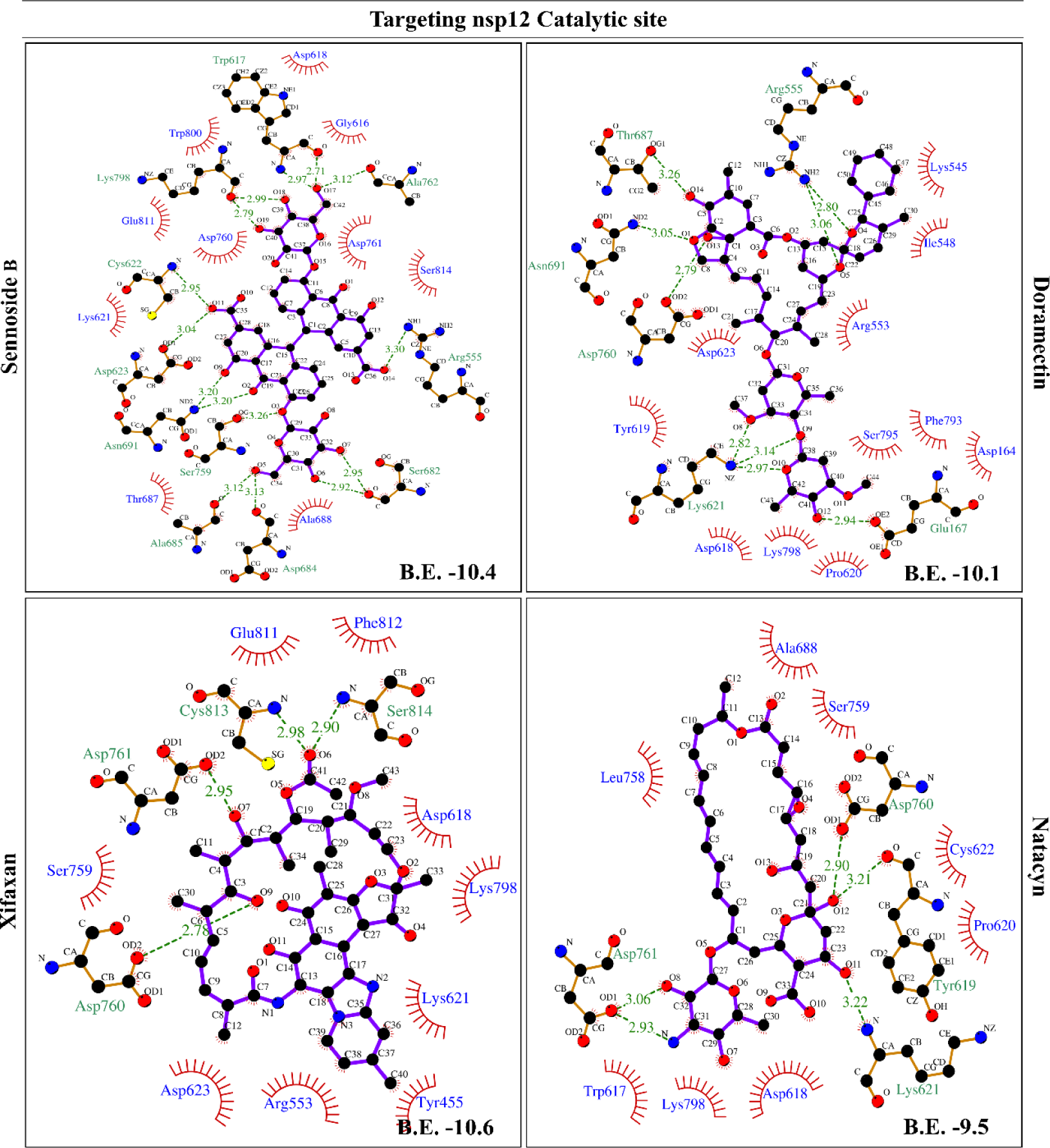
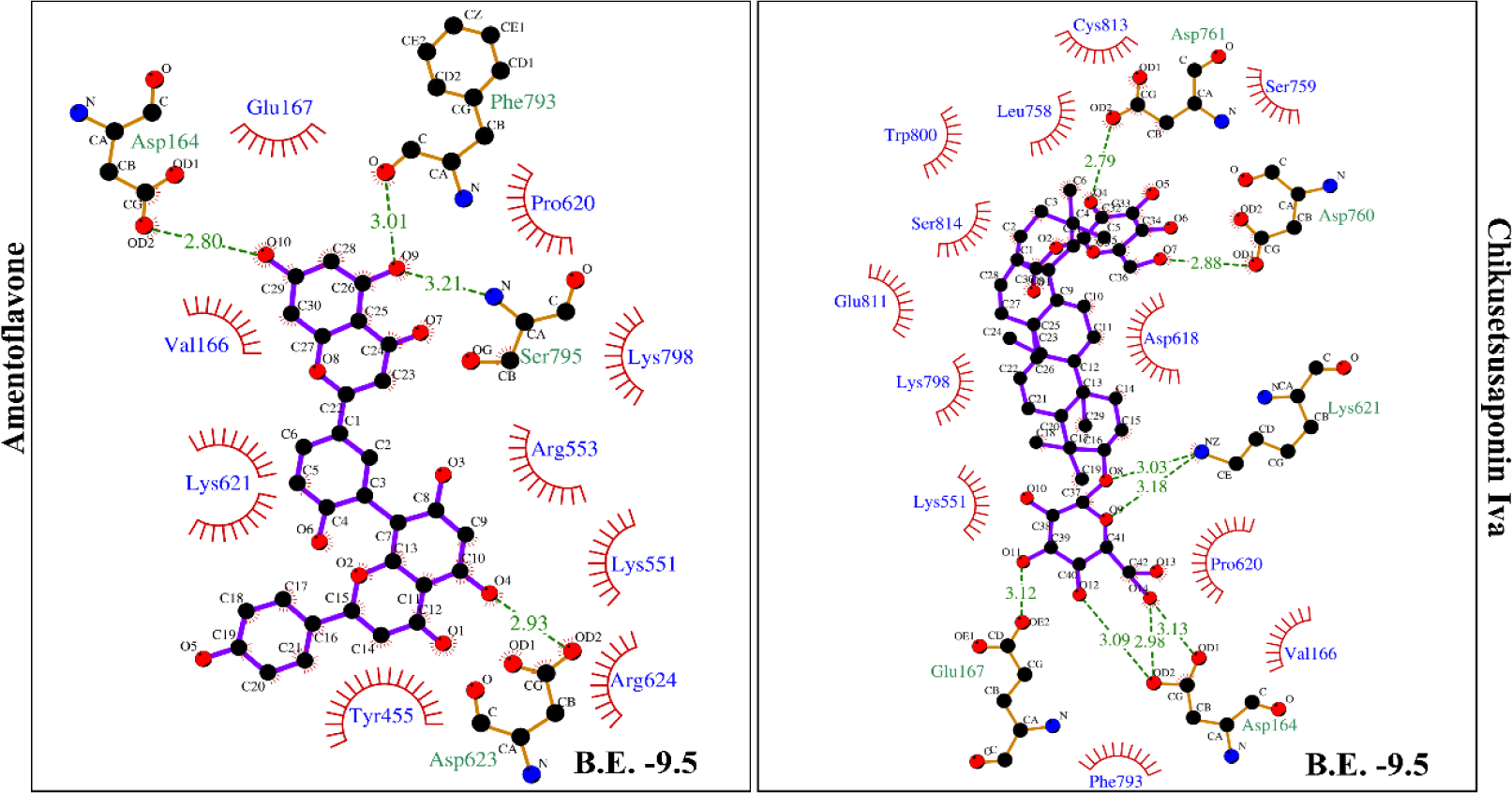
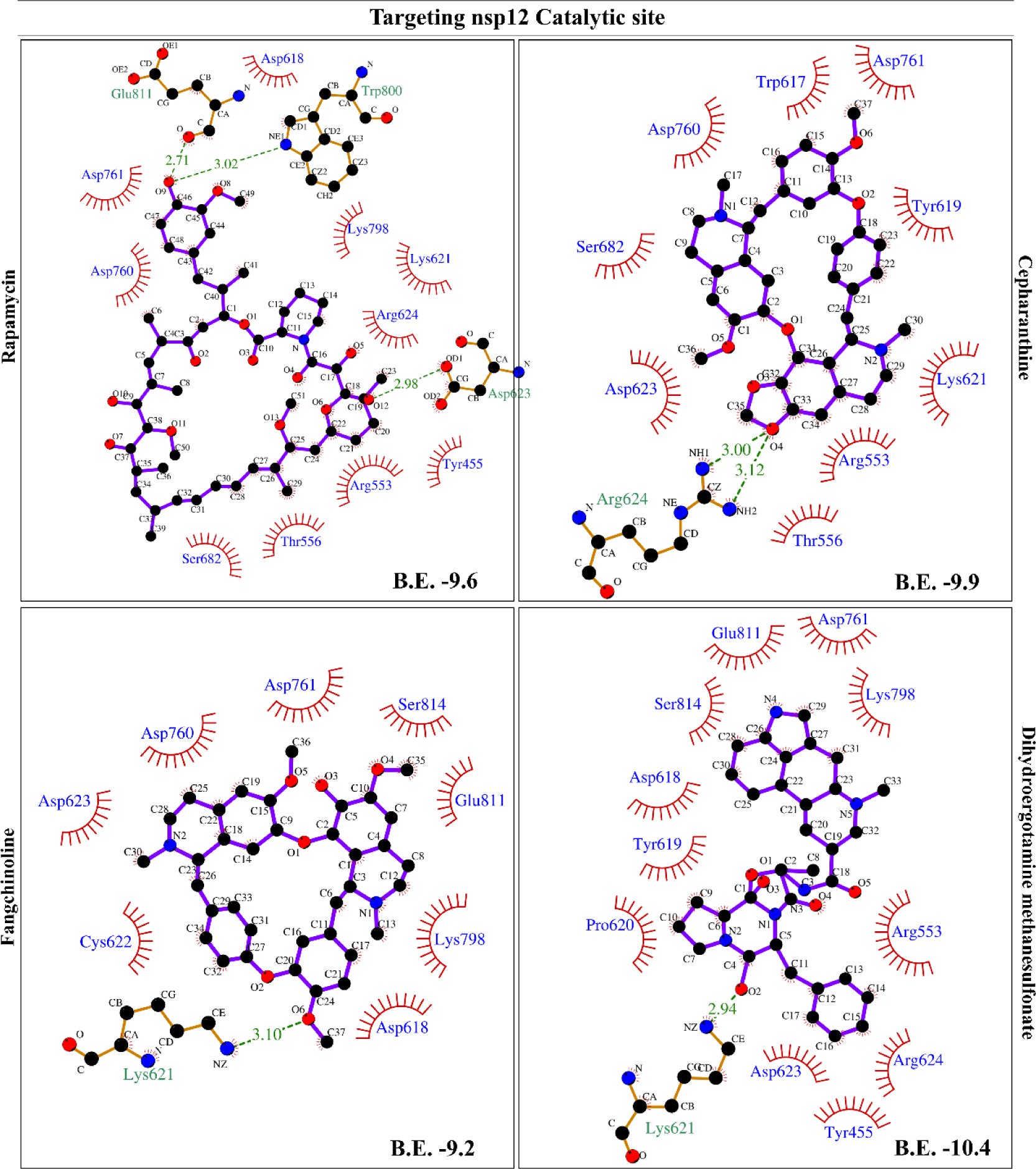
*Schematic presentation of sennoside B, doramectin, xifaxan, natacyn, amentoflavone, chikusetsusaponin iva, rapamycin, cepharanthine, fangchinoline, and dihydroergotamine methanesulfonate compounds at the catalytic site of nsp12 displaying H-bond, hydrophobic interactions and binding energy using the LIGPLOT+*.

**Figure 3:**
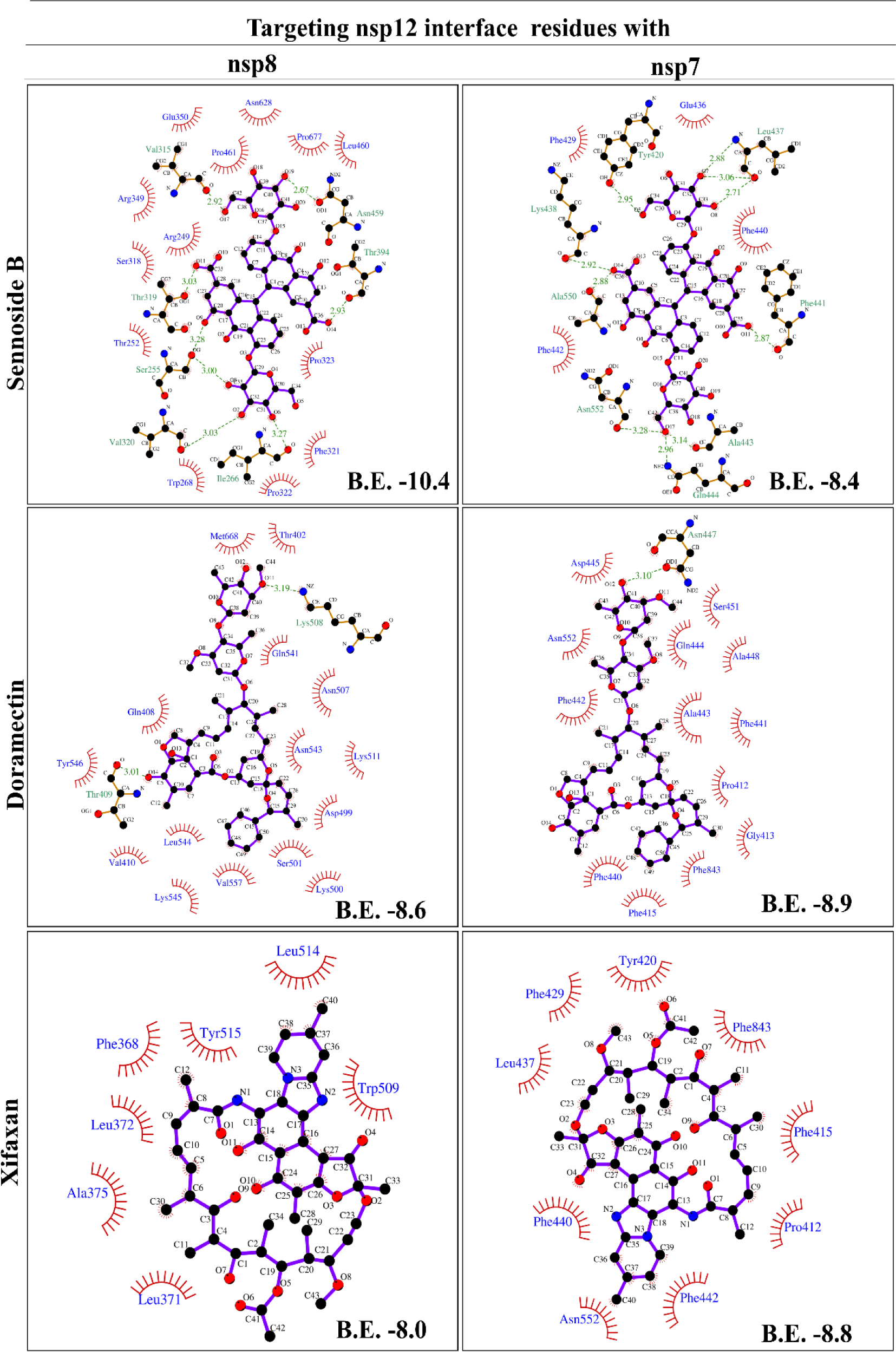
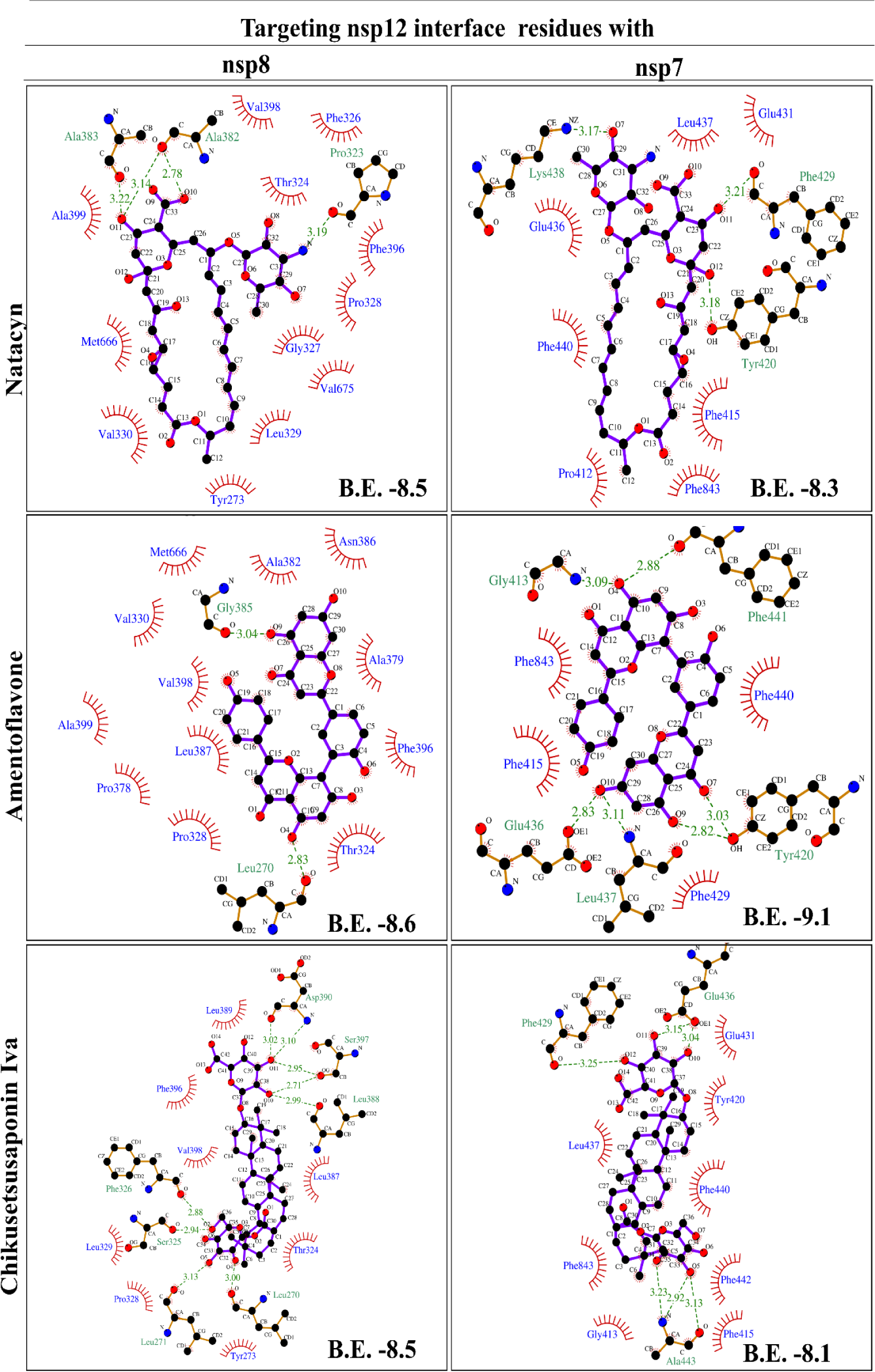
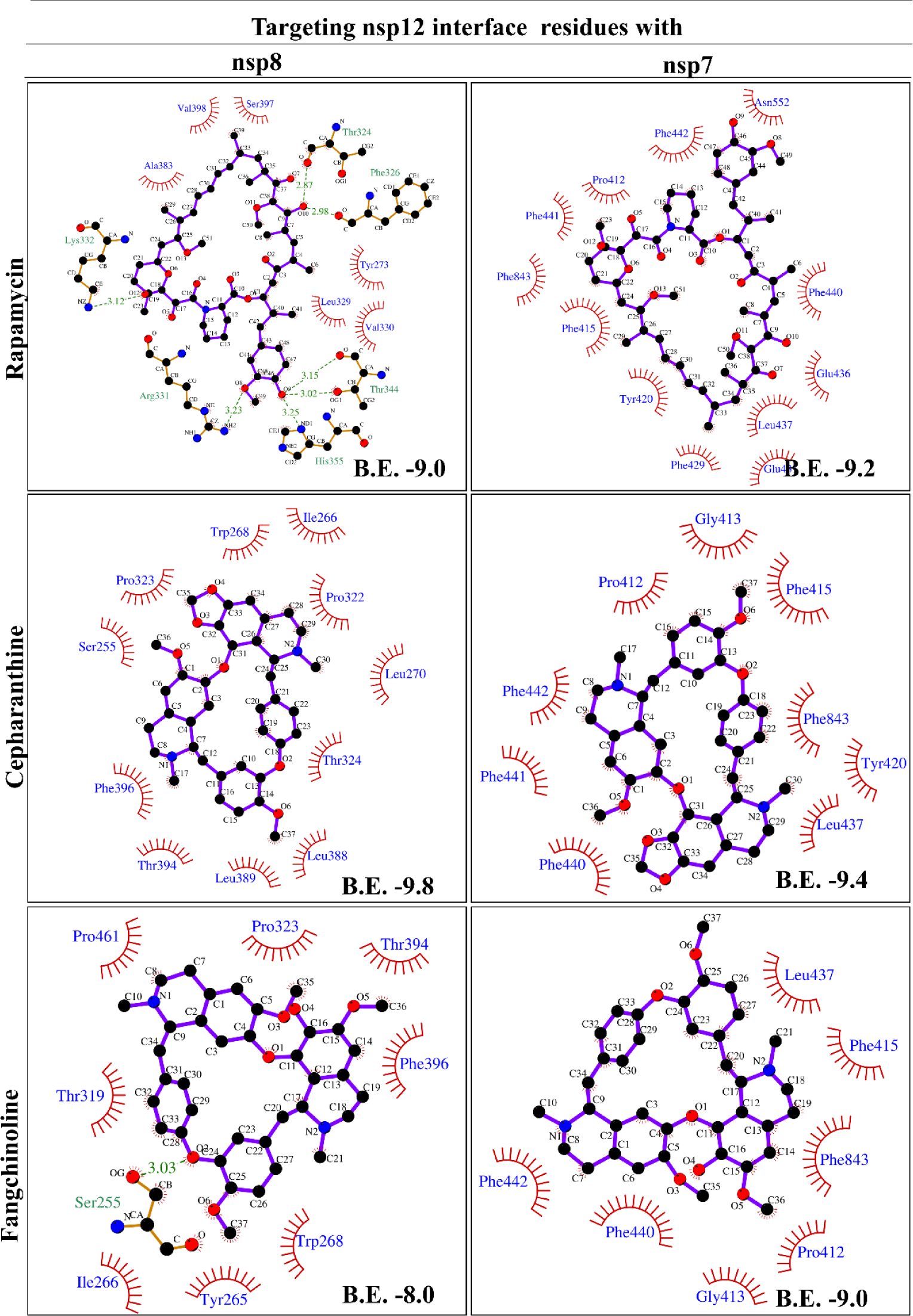
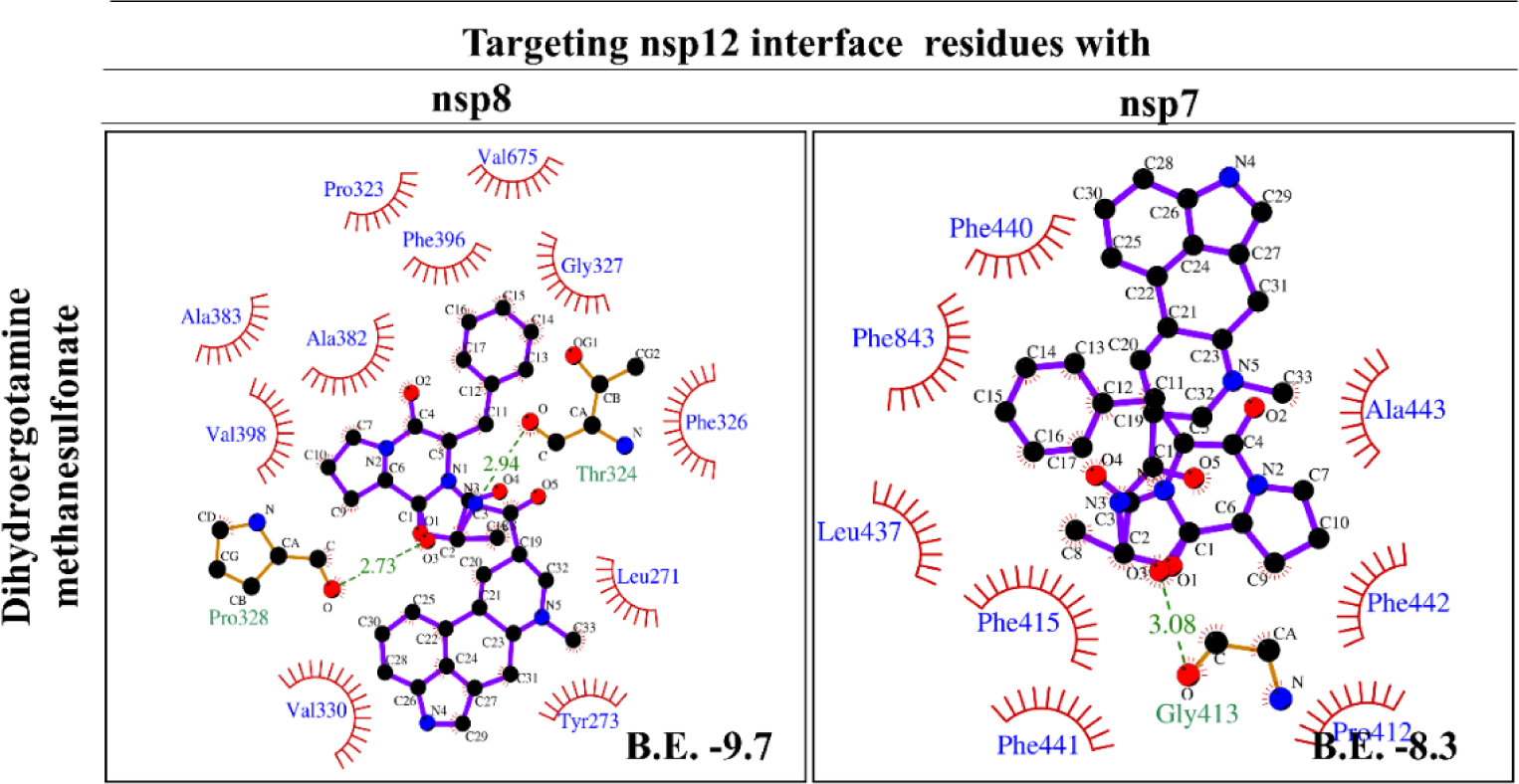
*Schematic presentation of sennoside B, doramectin, xifaxan, natacyn, amentoflavone, chikusetsusaponin iva, rapamycin, cepharanthine, fangchinoline, and dihydroergotamine methanesulfonate compounds at the interface site of nsp12 with nsp8 and nsp7 displaying H-bond, hydrophobic interactions and binding energy using the LIGPLOT+*.

**Figure 4:**
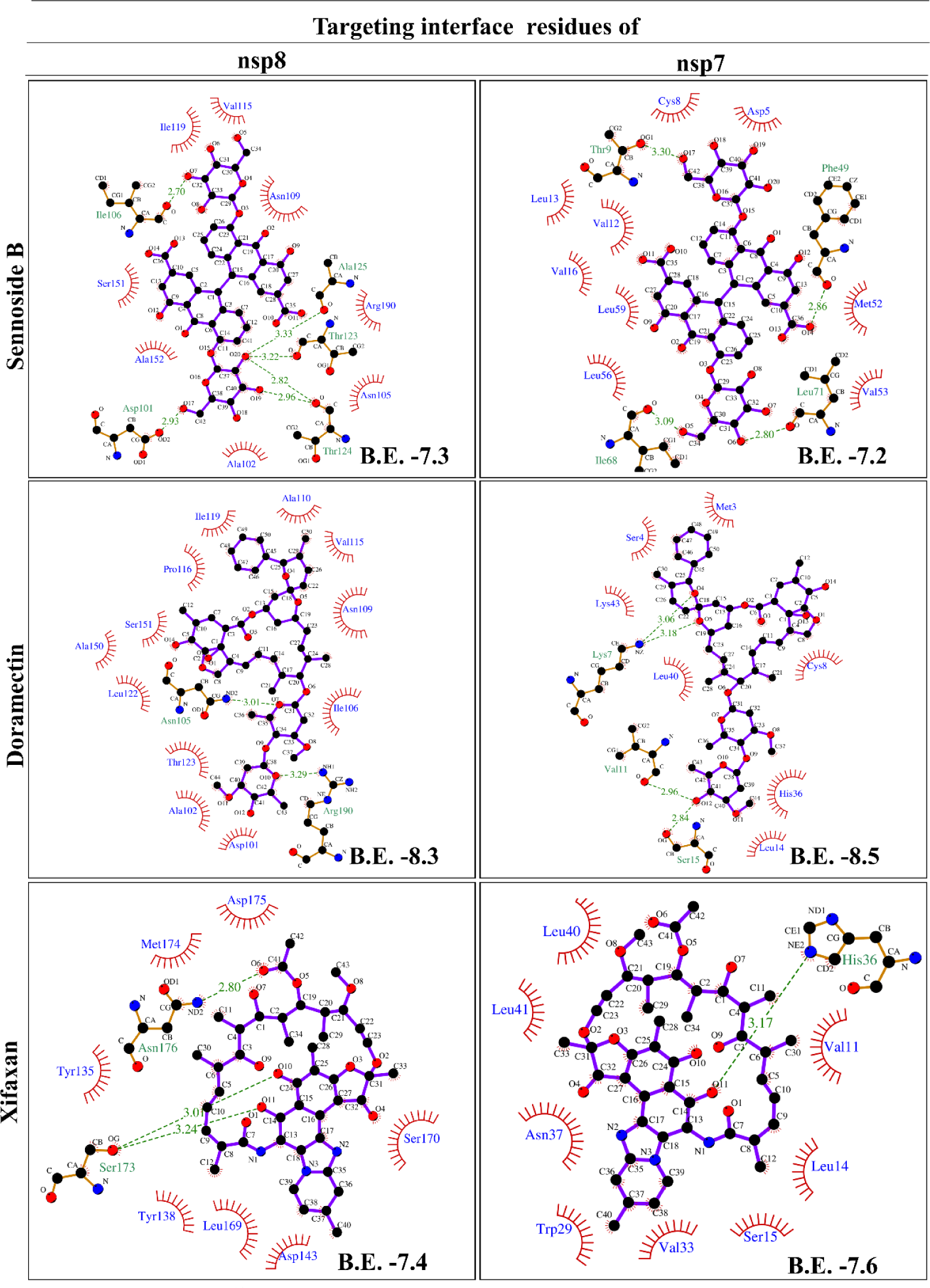
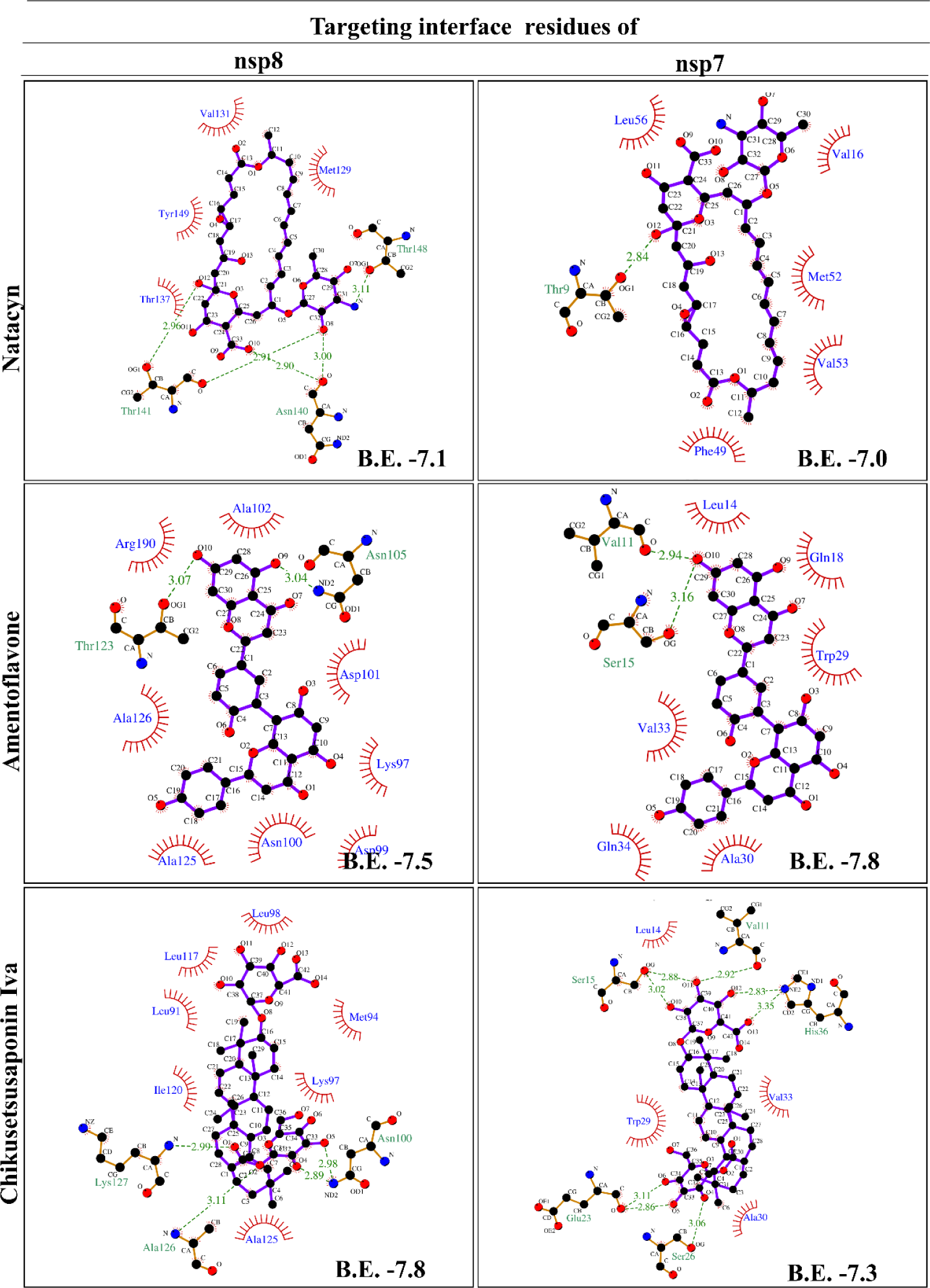
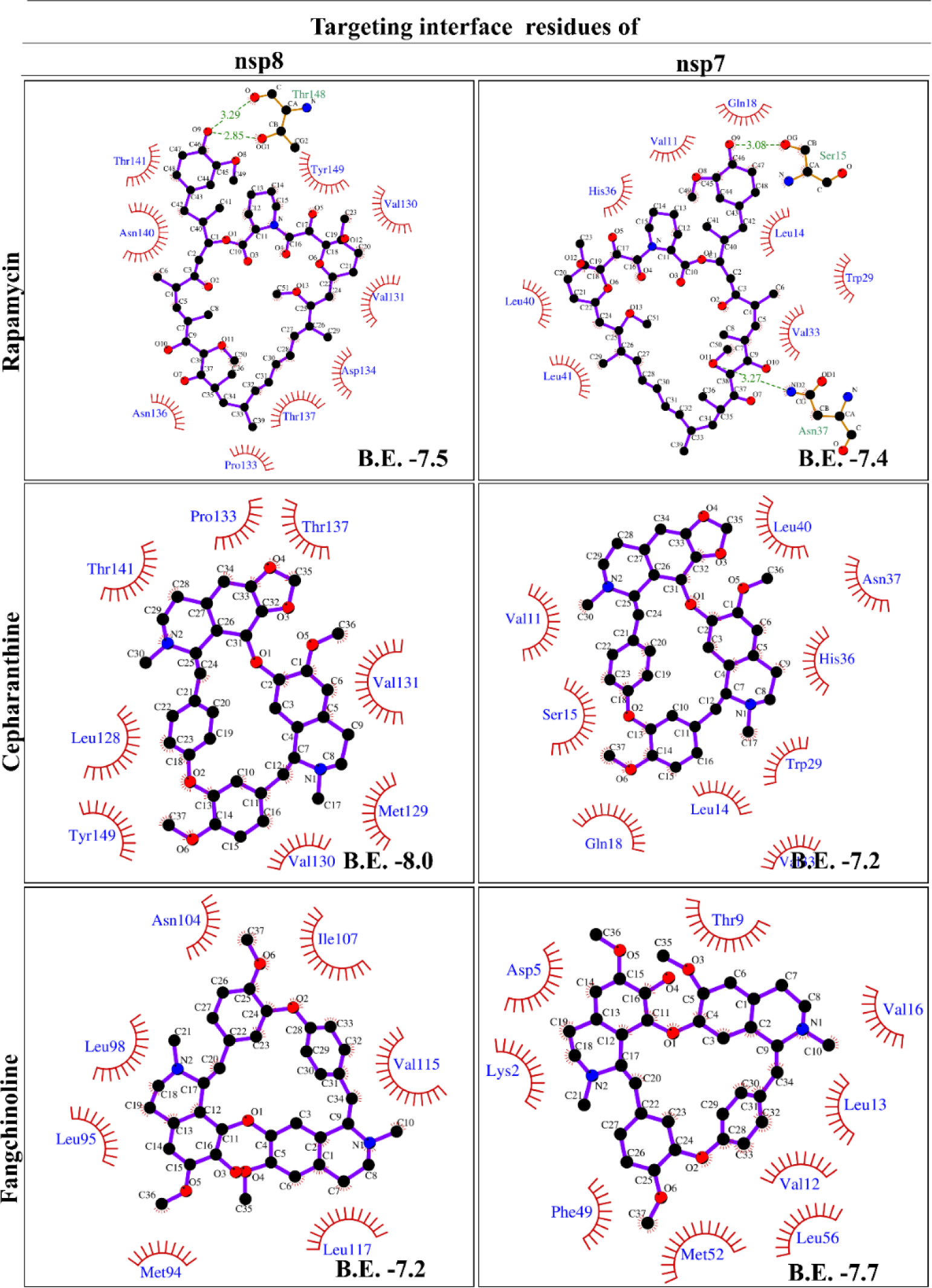
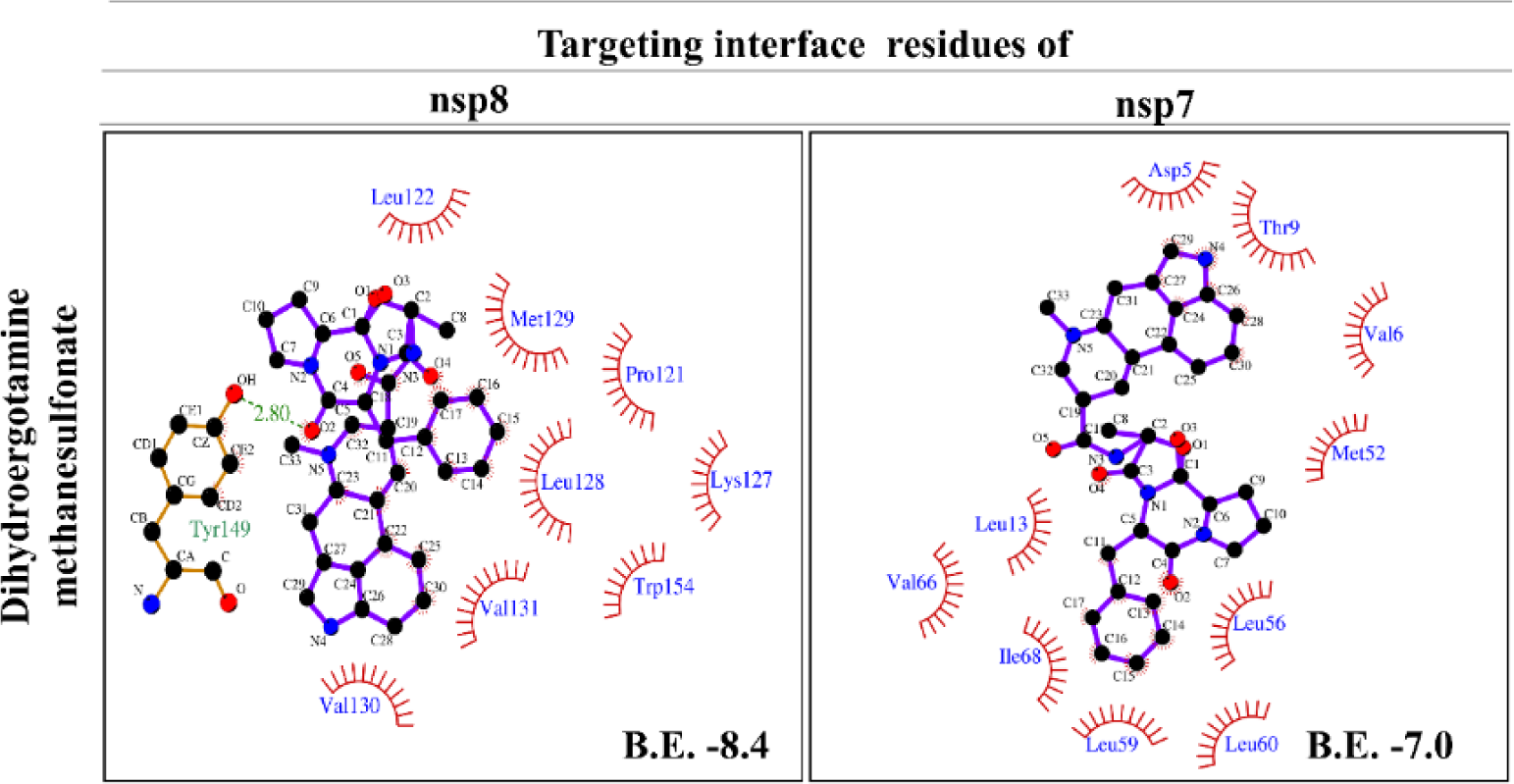
*Schematic presentation of sennoside B, doramectin, xifaxan, natacyn, amentoflavone, chikusetsusaponin iva, rapamycin, cepharanthine, fangchinoline, and dihydroergotamine methanesulfonate compounds at the interface site of nsp8 and nsp7 displaying H-bond, hydrophobic interactions and binding energy using the LIGPLOT+*.

**Table 2:** The hydrophobic interactions and H-bonds of identified compounds against the catalytic site, and the interface residues of SARS-CoV-2 nsp12 protein using the LIGPLOT+ analysis tool.

| Ligands |  | Catalytic<br>Vina | nsp12 interface residues against |  |
| --- | --- | --- | --- | --- |
|  |  |  | nsp8 | nsp7 |
| FDA Library from Selleckchem |  |  |  |  |
| Sennoside B | H bond (Å) | Arg555 (3.30), Tyr617 (2.71 & 2.97), Cys622 (2.95), Asp623 (3.04), Ser682 (2.92 & 2.95), Asp684 (3.13), Ala685 (3.12), Asn691 (3.20 & 3.20), Ser759 (3.26), Ala762 (3.12), Lys798 (2.79 & 2.99), | Ser255 (3.00 & 3.28), Ile266 (3.27), Val315 (2.92), Thr319 (3.03), Val320 (3.03), Thr394 (2.93), Asn459 (2.67) | Tyr420 (2.95), Leu437 (2.71, 2.88 & 3.06), Lys438 (2.92), Phe441 (2.87), Ala443 (3.14), Gln444 (2.96), Ala550 (2.88), Asn552 (3.28) |
|  | Hydrophobic | Gly616, Asp618, Lys621, Thr687, Ala688, Asp760, Asp761, Trp800, Glu811, Ser814 | Arg249, Thr252, Thr268, Ser318, Phe321, Pro322, Pro323, Arg349, Glu350, Leu460, Pro461, Asn628, Pro677 | Glu436, Phe429, Phe440, Phe442 |
| Doramectin | H bond (Å) | Glu167 (2.94), Arg555 (2.80 & 3.06), Lys621 (2.82, 2.97 & 3.14), Thr687 (3.26), Asn691 | Thr409 (3.01), Lys508 (3.19) | Asn447 (3.10) |
|  |  | (3.05), Asp760 (2.79), |  |  |
|  | Hydrophobic | Asp164, Ile548, Arg553, Lys545, Asp618, Tyr619, Pro620, Asp623, Phe793, Ser795, Lys798 | Thr402, Gln408, Val410, Asp499, Lys500, Ser501, Asn507, Lys511, Gln541, Asn543, Leu544, Lys545, Tyr546, Val557, Met668 | Pro412, Gly413, Phe415, Phe440, Phe441, Phe442, Ala443, Gln444, Asp445, Ala448, Ser451, Asn552, Phe843 |
| **Xifaxan** | H bond (Å) | Asp760 (2.89), Asp761 (3.01), Cys813 (3.04), Ser814 (2.86 & 2.98) | -- | -- |
|  | Hydrophobic | Tyr455, Arg553, Asp618, Asp623, Ser759, Lys798, Glu811, Phe812, | Leu371, Leu372, Phe368, Ala375, Trp509, Leu514, Tyr515 | Pro412, Phe415, Tyr420, Phe429, Leu437, Phe440, Phe442, Asn552, Phe843 |
| **Natural Product Library Compounds from Selleckchem** |  |  |  |  |
| **Amentoflavone** | H bond (Å) | Asp164 (2.80) Asp623 (2.93) Phe793 (3.01) Ser795(3.21) | Leu270 (2.83), Gly385 (3.04) | Gly413 (3.09), Tyr420 (2.82 & 3.03), Glu436 (2.83), Leu437 (3.11), Phe441 (2.88) |
|  | Hydrophobic | Val166, Tyr167, Tyr455, Arg551, Arg553, Pro620, Lys621, Arg624, Lys798, | Thr324, Pro328, Val330, Pro378, Ala379, Ala382, Asn386, Leu387, Phe396, Val398, Ala399, Met666 | Phe415, Phe429, Phe440, Phe843 |
| **Natacyn** | H bond (Å) | Tyr619 (3.21), Lys621 (3.22), Asp760 (2.90), Asp761 (2.93 & 3.06) | Pro323 (3.19), Ala382 (2.78 & 3.14), Ala383 (3.22) | Tyr420 (3.18), Phe429 (3.21), Lys438 (3.17) |
|  | Hydrophobic | Trp617, Pro620, Cys622, Asp618, Ala688, Lys758, Ser759, Lys798, | Tyr273, Thr324, Phe326, Gly327, Pro328, Leu329, Val330, Phe396, Val398, Ala399, Met666, Val675 | Glu436, Pro412, Phe415, Glu431, Leu437, Phe440, Phe843 |
| **Chikusetsusaponin Iva** | H bond (Å) | Asp164 (2.98, 3.13, 3.09), Glu167 (3.21), Lys621 (3.03 and 3.18), Asp760 (2.88), Asp761 (2.79), | Leu270 (3.00), Leu271 (3.13), Ser325 (2.94), Phe326 (2.88), Leu388 (2.99), Asp390 (3.20 & 3.10), Ser397 (2.71 & 2.95) | Phe429 (3.25), Glu436 (3.04 & 3.15), Ala443 (2.92, 3.13 & 3.23) |
|  | Hydrophobic | Val166, Lys551, Asp618, Phe620, Leu758, Ser759, Phe793, Lys798, Trp800, | Tyr273, Thr324, Pro328, Leu329, Leu387, Leu389, Phe396, Val398 | Gly413, Phe415, Leu437, Phe440, Tyr420, Glu431, Phe442, Phe843, |
|  |  | Glu811,<br>Cys813, Ser814 |  |  |
| **Rapamycin** | H bond (Å) | Asp623 (2.98),<br>Trp800 (3.02),<br>Glu811 (2.71) | Arg331 (3.23),<br>Lys332 (3.12),<br>Thr324 (2.87),<br>Phe326 (2.98),<br>Thr344 (3.02 &<br>3.15), His355 (3.25) | -- |
|  | Hydrophobic | Tyr455, Arg553,<br>Thr556,<br>Asp618,<br>Lys621,<br>Arg624, Ser682,<br>Asp760,<br>Asp761, Lys798 | Tyr273, Leu329,<br>Val330, Ala383,<br>Ser397, Val398 | Pro412, Phe415,<br>Tyr420, Phe429,<br>Glu436, Glu431,<br>Leu437, Phe440,<br>Phe441, Phe442,<br>Asn552, Phe843, |
| **Cepharanthine** | H bond (Å) | Arg624 (3.00<br>and 3.12) | -- | -- |
|  | Hydrophobic | Arg553, Thr556,<br>Trp617, Tyr619,<br>Lys621,<br>Asp623,<br>Asp760,<br>Asp761, Ser682 | Ser255, Ile266,<br>Trp268, Leu270,<br>Pro322, Pro323,<br>Thr324, Leu388,<br>Leu389, Thr394,<br>Phe396 | Pro412, Gly413,<br>Phe415, Tyr420,<br>Leu437, Phe440,<br>Phe441, Phe442,<br>Phe843 |
| **Fangchinoline** | H bond (Å) | Lys621 (3.10) | Ser255 (3.03) | -- |
|  | Hydrophobic | Asp618,<br>Cys622,<br>Asp623,<br>Asp760,<br>Asp761,<br>Lys798, Glu811,<br>Ser814, | Try265, Ile266,<br>Trp268, Thr319,<br>Pro323, Thr394,<br>Phe396, Pro461 | Pro412, Gly413,<br>Phe415, Leu437,<br>Phe440, Phe442,<br>Phe843 |
| **LOPAC library from Sigma** |  |  |  |  |
| **Dihydroergotamine**<br>**methanesulfonate** | H bond (Å) | Lys621 (2.94) | Thr324 (2.94),<br>Pro328 (2.73) | Gly413 (3.08) |
|  | Hydrophobic | Tyr455, Arg553,<br>Asp618,<br>Tyr619, Pro620,<br>Asp623,<br>Arg624,<br>Asp761,<br>Lys798, Glu811,<br>Ser814, | Leu271, Tyr273,<br>Pro323, Phe326,<br>Gly327, Val330,<br>Ala382, Ala383,<br>Phe396, Val398,<br>Val675 | Pro412, Phe415,<br>Leu437, Phe440,<br>Phe441, Phe442,<br>Ala443, Phe843, |

**Table 3:**
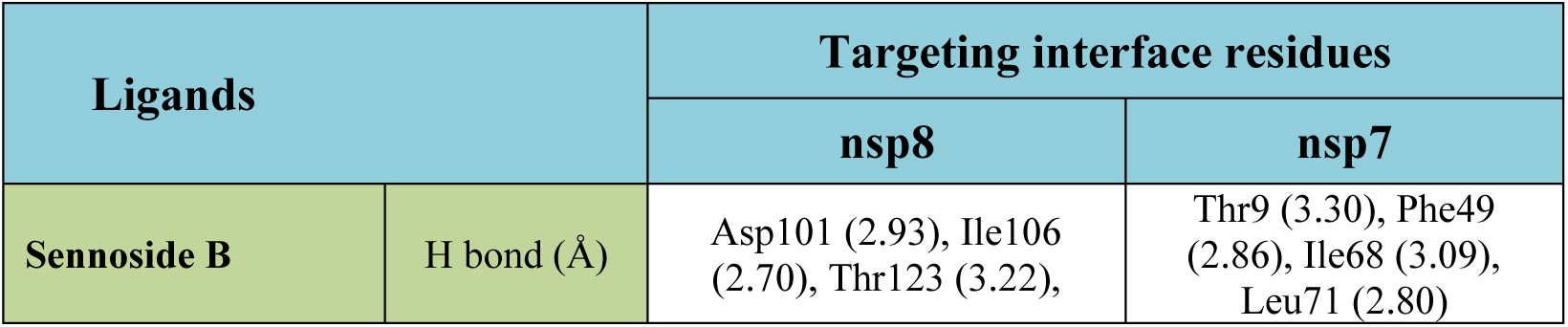
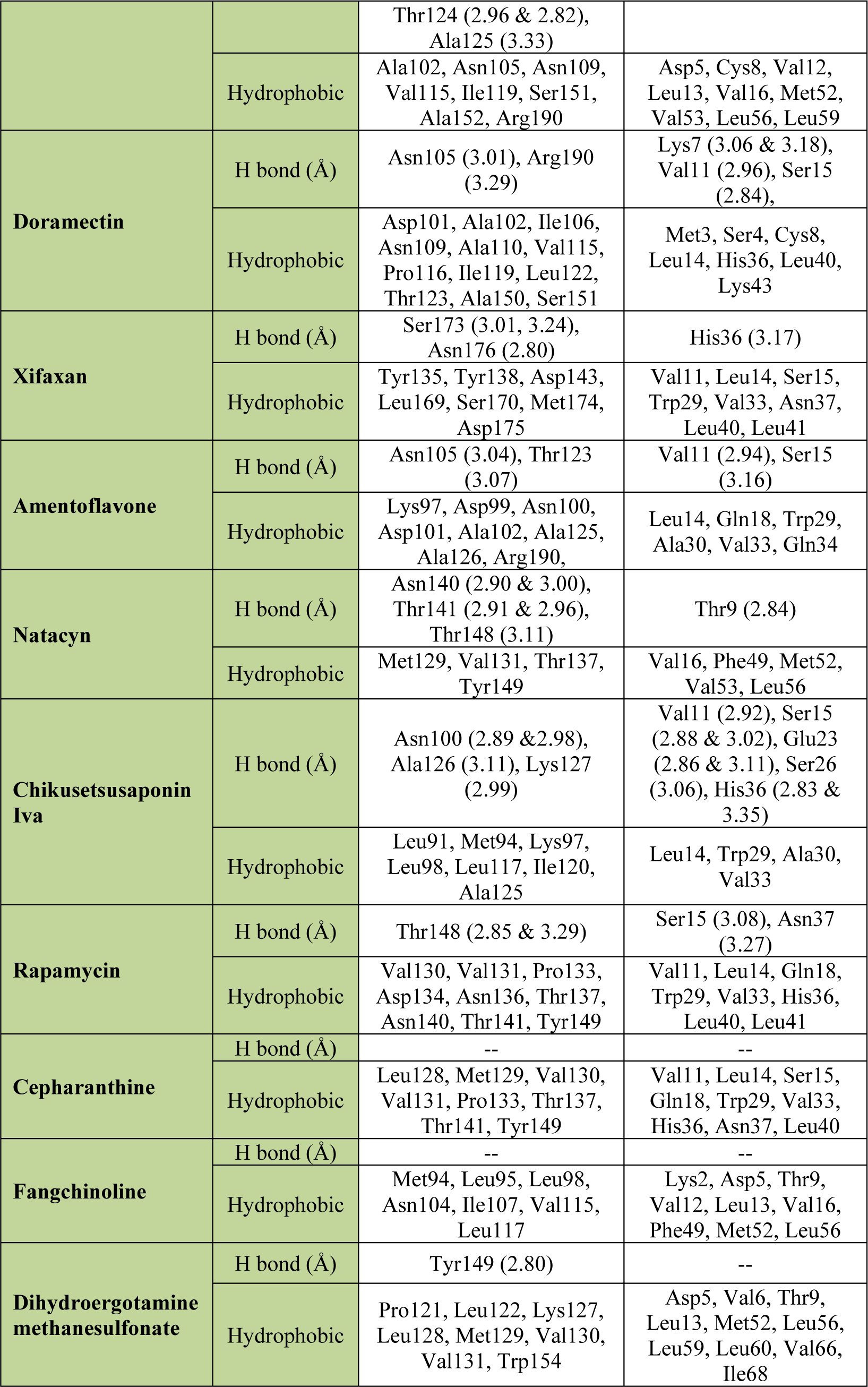
The hydrophobic interactions and H-bonds of identified compounds against the and the interface site of SARS-CoV-2 nsp7 and nsp8 protein.

| Ligands |  | Targeting interface residues |  |
| --- | --- | --- | --- |
|  |  | nsp8 | nsp7 |
| **Sennoside B** | H bond (Å) | Asp101 (2.93), Ile106 (2.70), Thr123 (3.22), | Thr9 (3.30), Phe49 (2.86), Ile68 (3.09), Leu71 (2.80) |
|  |  | Thr124 (2.96 & 2.82),<br>Ala125 (3.33) |  |
|  | Hydrophobic | Ala102, Asn105, Asn109,<br>Val115, Ile119, Ser151,<br>Ala152, Arg190 | Asp5, Cys8, Val12,<br>Leu13, Val16, Met52,<br>Val53, Leu56, Leu59 |
| **Doramectin** | H bond (Å) | Asn105 (3.01), Arg190<br>(3.29) | Lys7 (3.06 & 3.18),<br>Val11 (2.96), Ser15<br>(2.84), |
|  | Hydrophobic | Asp101, Ala102, Ile106,<br>Asn109, Ala110, Val115,<br>Pro116, Ile119, Leu122,<br>Thr123, Ala150, Ser151 | Met3, Ser4, Cys8,<br>Leu14, His36, Leu40,<br>Lys43 |
| **Xifaxan** | H bond (Å) | Ser173 (3.01, 3.24),<br>Asn176 (2.80) | His36 (3.17) |
|  | Hydrophobic | Tyr135, Tyr138, Asp143,<br>Leu169, Ser170, Met174,<br>Asp175 | Val11, Leu14, Ser15,<br>Trp29, Val33, Asn37,<br>Leu40, Leu41 |
| **Amentoflavone** | H bond (Å) | Asn105 (3.04), Thr123<br>(3.07) | Val11 (2.94), Ser15<br>(3.16) |
|  | Hydrophobic | Lys97, Asp99, Asn100,<br>Asp101, Ala102, Ala125,<br>Ala126, Arg190, | Leu14, Gln18, Trp29,<br>Ala30, Val33, Gln34 |
| **Natacyn** | H bond (Å) | Asn140 (2.90 & 3.00),<br>Thr141 (2.91 & 2.96),<br>Thr148 (3.11) | Thr9 (2.84) |
|  | Hydrophobic | Met129, Val131, Thr137,<br>Tyr149 | Val16, Phe49, Met52,<br>Val53, Leu56 |
| **Chikusetsusaponin**<br>**Iva** | H bond (Å) | Asn100 (2.89 & 2.98),<br>Ala126 (3.11), Lys127<br>(2.99) | Val11 (2.92), Ser15<br>(2.88 & 3.02), Glu23<br>(2.86 & 3.11), Ser26<br>(3.06), His36 (2.83 &<br>3.35) |
|  | Hydrophobic | Leu91, Met94, Lys97,<br>Leu98, Leu117, Ile120,<br>Ala125 | Leu14, Trp29, Ala30,<br>Val33 |
| **Rapamycin** | H bond (Å) | Thr148 (2.85 & 3.29) | Ser15 (3.08), Asn37<br>(3.27) |
|  | Hydrophobic | Val130, Val131, Pro133,<br>Asp134, Asn136, Thr137,<br>Asn140, Thr141, Tyr149 | Val11, Leu14, Gln18,<br>Trp29, Val33, His36,<br>Leu40, Leu41 |
| **Cepharanthine** | H bond (Å) | -- | -- |
|  | Hydrophobic | Leu128, Met129, Val130,<br>Val131, Pro133, Thr137,<br>Thr141, Tyr149 | Val11, Leu14, Ser15,<br>Gln18, Trp29, Val33,<br>His36, Asn37, Leu40 |
| **Fangchinoline** | H bond (Å) | -- | -- |
|  | Hydrophobic | Met94, Leu95, Leu98,<br>Asn104, Ile107, Val115,<br>Leu117 | Lys2, Asp5, Thr9,<br>Val12, Leu13, Val16,<br>Phe49, Met52, Leu56 |
| **Dihydroergotamine**<br>**methanesulfonate** | H bond (Å) | Tyr149 (2.80) | -- |
|  | Hydrophobic | Pro121, Leu122, Lys127,<br>Leu128, Met129, Val130,<br>Val131, Trp154 | Asp5, Val6, Thr9,<br>Leu13, Met52, Leu56,<br>Leu59, Leu60, Val66,<br>Ile68 |

Moreover, the LIGPLOT^+^ analysis of all ligands revealed the H bond and hydrophobic interactions between the ligands and SARS-CoV-2 proteins (**Figure 2-4**). Based on the interaction, sennoside B exhibits the highest B.E. (kcal/mol) with the highest number of H bond when compared with different ligands for different sites (**Figure 2-4** and **Table 2-3**). While doramectin, cepharanthine, and dihydroergotamine methanesulfonate shows the highest hydrophobic bonds with the catalytic site of nsp12 as well as with the interface residues of nsp12, nsp7 and nsp8 (**Figure 2-4** and **Table 2-3**). Moreover, the complete analysis of H bond and hydrophobic bond interaction of compounds are presented in (**Figure 2-4** and **Table 2-3**).

### Expression and purification of nsp8 and nsp7 recombinant proteins

The gene sequence of nsp8 (1-198 amino acid residues) and nsp7 (1-84 amino acid residues) were cloned into an expression vector pET28c carrying a 6x-histidine affinity tag. The DH5α *E. Coli* cells were transformed, and the plasmid was isolated to validate the transformed DH5α *E. Coli* cells using Sanger sequencing. The recombinant proteins were expressed in *E. coli* BL21 (DE3) cells and purified using immobilized Ni^+2^-NTA affinity chromatography. The eluted fractions were resolved on 12% SDS-PAGE. The Coomassie brilliant blue stained lanes of elution fractions displayed bands of molecular weight ∼24 kDa, and ∼10.4 kDa, corresponding to 6xHis-nsp8 and 6xHis-nsp7 protein, respectively (**Supplementary figure 2**).

### Binding kinetics of identified compounds

SPR experiments with a Biacore T200 system (Biacore Inc., Uppsala, Sweden) were performed to determine the binding affinities and kinetics of the selected compounds. The SPR technique is highly sensitive, enabling real-time detection of direct binding interaction. It provides three essential parameters, dissociation rate (*k*_off_), association rate (*k*_on_), and binding affinity (*K*D) at equilibrium. We monitored the binding behaviours of the ten potential compounds to his-tagged nsp7 and nsp8 proteins of SARS-CoV-2 at progressively increasing concentrations to observe the dose-response. As shown in **Figure 5-6**, all selected compounds exhibited concentration-dependent binding to the nsp8 and nsp7 proteins of the SARS-CoV-2. All these compounds showed stronger binding affinities, with a dissociation constant K_D_ value in the micromolar range except natacyn for nsp7 protein. The binding kinetics data of these compounds are shown in **Table 4** and **Figures 5-6**.

**Figure 5:**
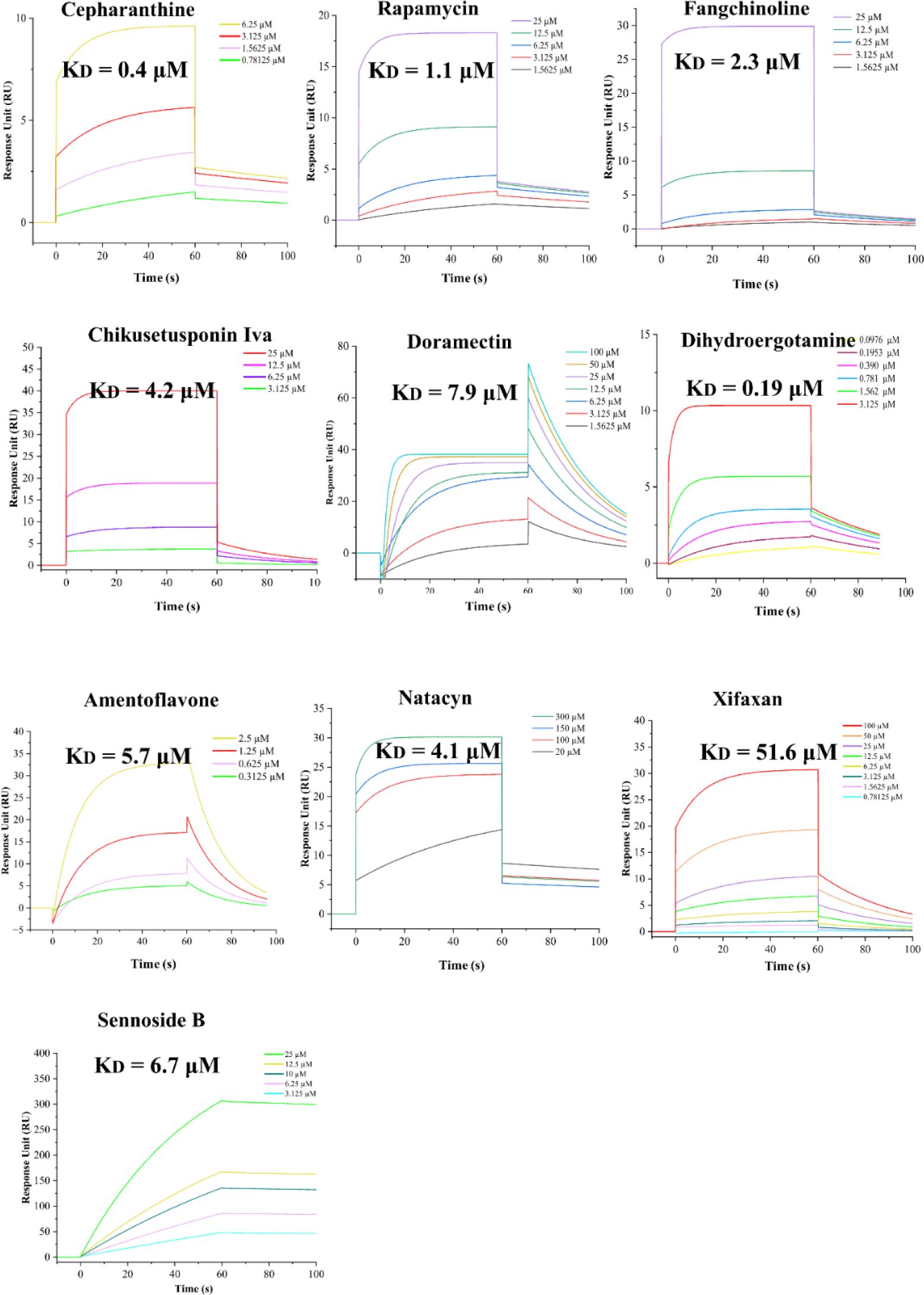
*Biacore T200 was used to perform binding kinetics and affinity analyses on all selected drugs for binding to the nsp8 protein of SARS-CoV-2. Different concentrations of the compounds were tried into the SPR Biacore T200 machine to fit the kinetic data, which are represented by different colors in each graph*.

**Figure 6:**
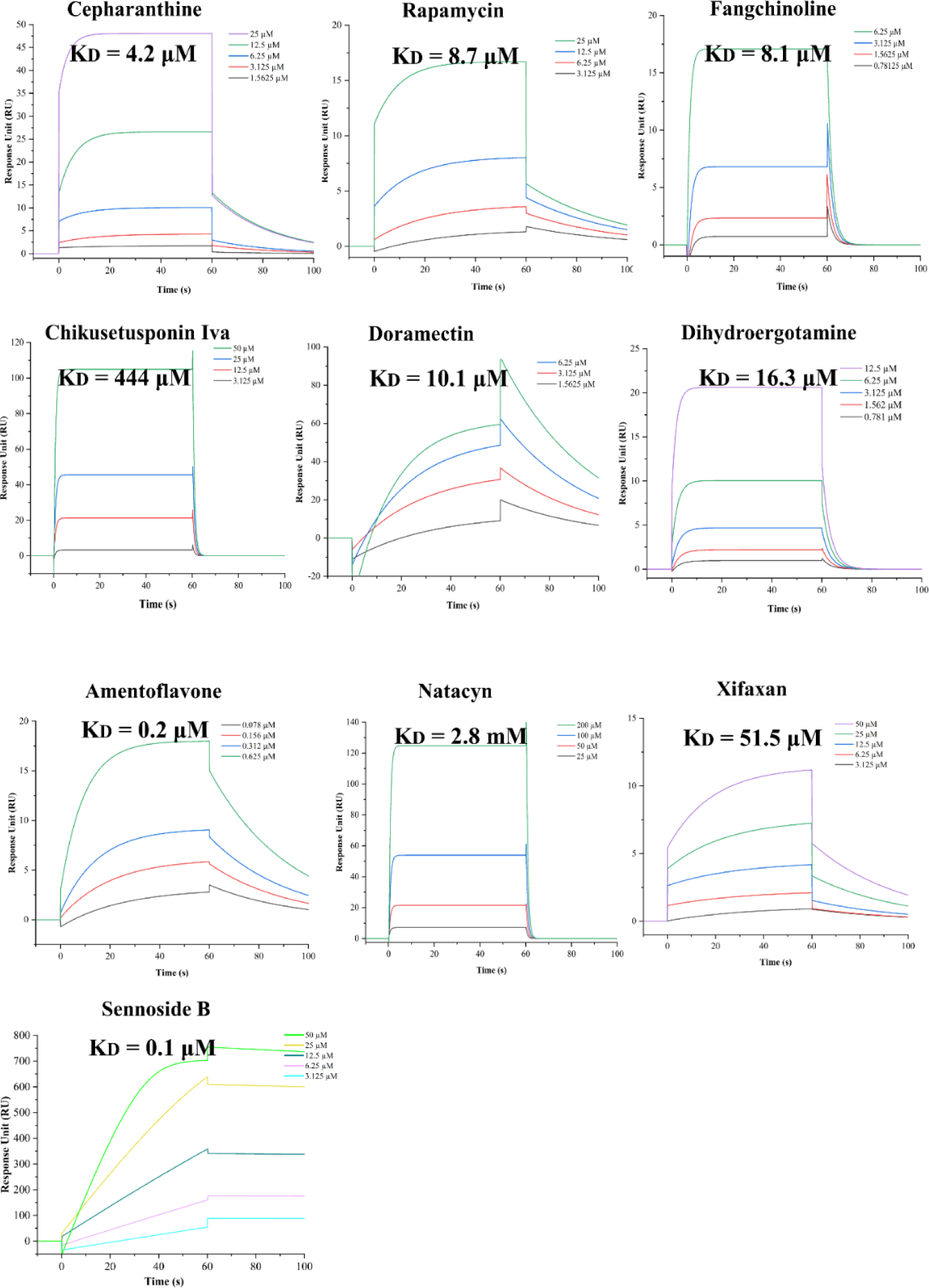
*Biacore T200 was used to perform binding kinetics and affinity analyses on all selected drugs for binding to the nsp7 protein of SARS-CoV-2. Different concentrations of the compounds were tried in the SPR Biacore T200 machine to fit the kinetic data, represented by different colors in each graph*.

**Table 4:** Binding kinetics and affinity analysis of all selected compounds for binding to the nsp8 and nsp7 protein of SARS-CoV-2 using a Biacore T200.

| Compounds | Biophysical Data |  |  |  |  |  |
| --- | --- | --- | --- | --- | --- | --- |
|  | SARS-CoV-2 nsp8 |  |  | SARS-CoV-2 nsp7 |  |  |
| | $K_D$<br>($\mu\text{M}$) | $k_a$ ($\text{M}^{-1} \cdot \text{s}^{-1}$) | $k_d$ (1/s) | $K_D$<br>($\mu\text{M}$) | $k_a$ ($\text{M}^{-1} \cdot \text{s}^{-1}$) | $k_d$ (1/s) |
| **Cepharanthine** | 0.40 | 13750 | 0.005608 | 4.28 | 9806 | 0.04198 |
| **Rapamycin** | 1.12 | 7009 | 0.007863 | 8.76 | 3077 | 0.02695 |
| **Fangchinoline** | 2.30 | 6849 | 0.01581 | 8.14 | 58410 | 0.4757 |
| **Chikusetsusaponin**<br>**Iva** | 4.21 | 8017 | 0.03378 | 444 | 2998 | 1.331 |
| **Doramectin** | 7.95 | 4976 | 0.03958 | 10.1 | 2733 | 0.02762 |
| **Dihydroergotamine**<br>**methanesulfonate** | 0.19 | 117700 | 0.02269 | 16.3 | 19290 | 0.3154 |
| **Amentoflavone** | 5.79 | 11310 | 0.06553 | 0.25 | 120300 | 0.03072 |
| **Natacyn** | 4.11 | 786.6 | 0.003235 | 2865 | 514.3 | 1.474 |
| **Xifaxan** | 51.6 | 585.9 | 0.03024 | 51.5 | 531.1 | 0.02736 |
| **Sennoside B** | 0.67 | 787.9 | 52950 | 0.18 | 3629 | 0.0006719 |

**Table 5:** Reported roles of identified antiviral compounds.

| Ligands | Functions | Reference |
| --- | --- | --- |
| **FDA Library from Selleckchem** |  |  |
| **Sennoside B** | Treat constipation and empty the large intestine before surgery | (Dreessen et al., 1981; Esposito et al., 2016; Hardcastle and Wilkins, 1970; Kon et al., 2014) |
| **Doramectin** | Antiparasitic agent | (Dreessen et al., 1981) |
| **Xifaxan** | Treat diarrhea | (Rifaximin, 2018) |
| **Natural Product Library Compounds from Selleckchem** |  |  |
| **Amentoflavone** | Anti-inflammatory, anxiolytic, anti-oxidative, anti-cancer, and neuroprotective effects | (Beaulieu et al., 1996; Hanioka et al., 2008; Jenkinson et al., 2012; Liu et al., 2016) |
| **Natacyn** | Treat the symptoms of fungal keratitis, and fungal blepharitis or conjunctivitis | (Arora et al., 2011; He et al., 2019; Raab, 1972) |
| **Chikusetsusaponin Iva** | Anti-inflammatory mediators, antiviral, and antithrombotic effect | (Dahmer et al., 2012; Rattanathongkom et al., 2009; Xu et al., 2021) |
| **Rapamycin** | Immunosuppressive, antitumor, neuroregenerative or neuroprotective, inhibits mammalian target of rapamycin (mTOR) | (Houchens et al., 1983; Malagelada et al., 2010; Martel et al., 1977; Pan et al., 2008; Tain et al., 2009) |
| **Cepharanthine** | Anti-oxidative properties, anti-inflammatory properties, and anti-lipid peroxidation activity | (Halicka et al., 2008; Kogure et al., 2003; Samra et al., 2016) |
| **Fangchinoline** | Anti- HIV-1 inhibitor | (Wan et al., 2012) |
| **LOPAC library from Sigma** |  |  |
| **Dihydroergotamine methanesulfonate** | Antimigraine characteristics | (Villalon et al., 1992) |

### Antiviral efficacy against SARS-CoV-2

The percent viability for various compounds on Vero cells was determined, revealing their effects on cell viability after 48 h. All selected compounds demonstrated over 70% viability at ∼100 μM, except for cepharanthine, fangchinoline, and doramectin. These three compounds exhibited over 70% viability at lower concentrations, starting from 25 μM for cepharanthine and fangchinoline and 6.25 μM for doramectin, confirming their non-toxic nature in this concentration range (**Supplementary figure 3**). Therefore, the maximum concentration for the antiviral experiments was set at a threshold viability of >80% to ensure a safe therapeutic window between effective antiviral activity and cell toxicity.

Compounds potency against SARS-CoV-2 *in vitro* has been studied in Vero cells for 48 h using qRT-PCR. The results demonstrated that the virus titre in Vero cells treated with range of concentration for amentoflavone, xifaxan, natacyn, sennoside B, cepharanthine, fangchinoline, and doramectin was reduced significantly compared to the virus control (VC) (**Figure 7**). At a very low concentration, namely 6 μM, cepharanthine, fangchinoline, and doramectin exhibited a notable reduction of ∼99% in the relative RNA levels of SARS-CoV-2 (**Figure 7**). This reduction is highly significant for all these selected compounds. Moreover, amentoflavone, xifaxan, natacyn, and sennoside B also demonstrated dose-dependent antiviral potential against SARS-CoV-2 in Vero cells, starting from their highest viability concentration range.

**Figure 7:**
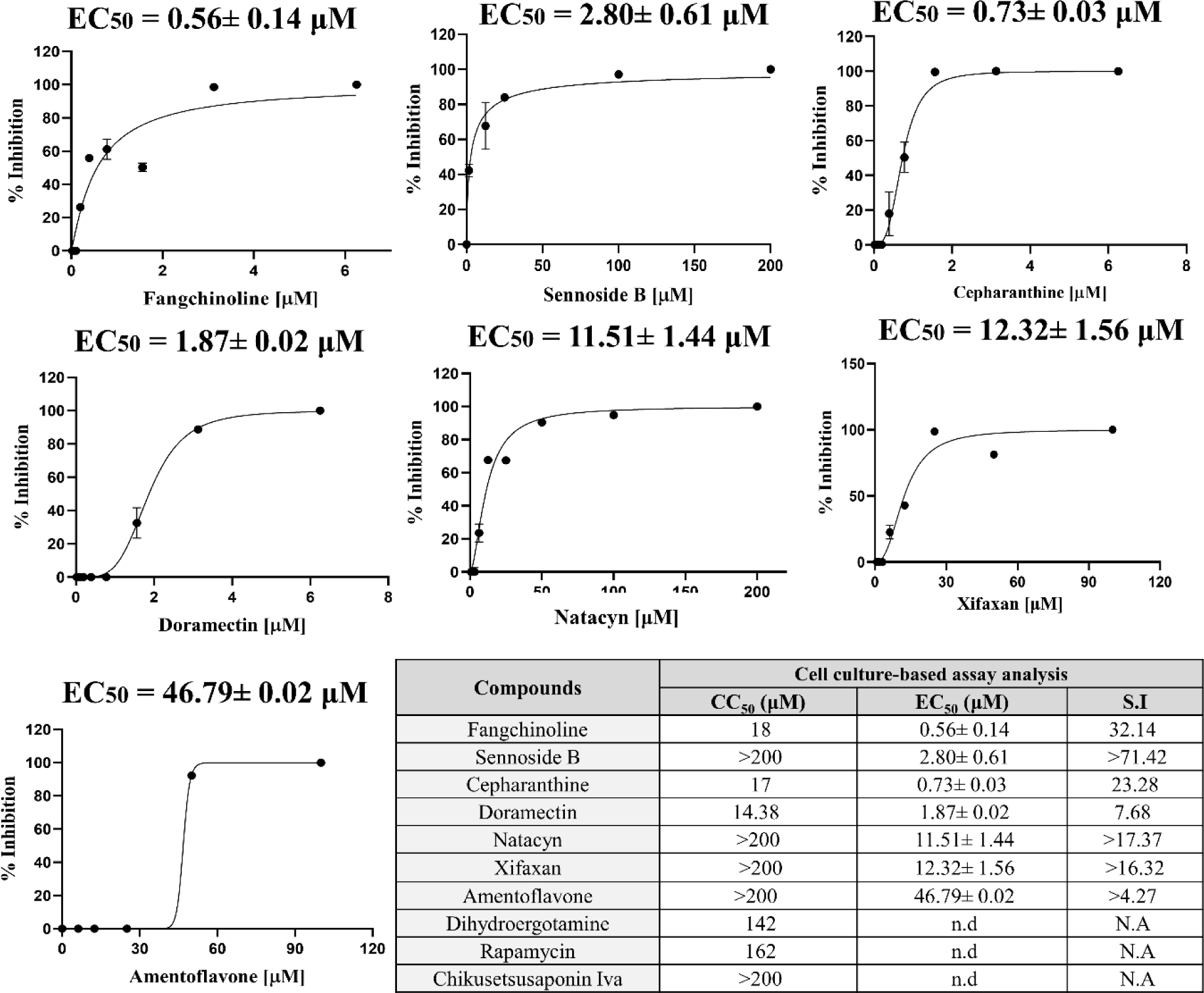
*Inhibiton assay of selected compounds in Vero cells. Vero cells were exposed to a series of two-fold diluted concentrations of the compounds and incubated at 37 °C in 5% CO_2_ for 48 h. The chosen range of cytotoxic data, where cell viability exceeded 80%, indicates a favorable cytotoxic profile for evaluating the antiviral potency against SARS-CoV-2. The relative activity of infected Vero cells was considered as virus control, and the percentage inhibition resulting from compound treatment is expressed as the activity inhibition relative to the infected cells left untreated. Each data point represents three independent replicate experiments. (N.A specifies not applicable and n.d denotes not determined)*.

To further evaluate the antiviral activity, we determined the EC_50_ and selectivity index (S.I.) of all these compounds against SARS-CoV-2 (Figure 8). Fangchinoline exhibited the lowest EC_50_ value of 0.56 µM, while other compounds gave the EC_50_ values in the micromolar range, typically less than 50 µM. Based on the S.I, we further selected fangchinoline and sennoside B compounds for *in vivo* testing against SARS-CoV-2, as they exhibited a highly favorable S.I (**Table shown in Figure 7**). However, dihydroergotamine methanesulfonate, chikusetsusaponin iva, and rapamycin showed no activity against SARS-CoV-2, as observed in our study.

To corroborate the findings obtained from the qRT-PCR results, the inhibition of virus replication in the presence of increasing concentration of drug was also evaluated by TCID_50_ assay. The cell lysate infected and compound treated cells was diluted ten-fold ranging from 10^1^ to 10^8^ TCID_50_ and was incubated with Vero cell monolayer. Similarly, virus control (Infected and compound non-treated cells) cell lysate was also processed for TCID_50_. In accordance with qRT-PCR results, effective compounds treatement resulted into decreased virus titre. Control monolayers infected with SARS-CoV-2 exhibited CPE, while treated infected cells demonstrated inhibitory effects against the virus at similar doses as tested in RT-PCR. The virus titration assay revealed a concentration-dependent reduction in viral titer upon treatment with the compounds.

Notably, at highest tested concentration, a significant reduction in viral titer was observed compared to the untreated infected positive control for cepharanthine (6 log TCID_50_/mL decrease), fangchinoline (5 log TCID_50_/mL decrease), and doramectin (6 log TCID_50_/mL decrease), sennoside B (3.5 log TCID_50_/mL decrease), amentoflavone (4 log TCID_50_/mL decrease), natacyn (4.5 log TCID_50_/mL decrease), xifaxan (2.5 log TCID_50_/mL decrease) (**Supplementary figure 4**). Conversely, dihydroergotamine methanesulfonate, chikusetsusaponin iva, and rapamycin exhibited no inhibitory potential against SARS-CoV-2. These results aligned with the qRT-PCR findings and confirmed that all seven potential compounds effectively inhibit viral replication in SARS-CoV-2-infected cells, ultimately protecting cellular monolayers against viral infections.

## Discussion

Since the beginning of the pandemic, a significant number of investigations and clinical trials have been initiated, reflecting the urgent need to identify effective antiviral treatments amidst the global burden of new strains of SARS-CoV-2. Therefore, extensive research is ongoing to screen various compounds targeting different core conserved SARS-CoV-2 proteins, including small molecules, antiviral peptides, nucleoside analogs, natural plant metabolites, etc. The primary objective of all these studies is to identify the most potent antiviral molecules that can be universally employed for the treatment of SARS-CoV-2, offering hope for effective therapeutic interventions against the virus. However, developing a novel treatment for COVID-19 in a short period will be difficult. One efficient method is to identify potential compounds by employing repurposing approach.

In this study, we employed a repurposing approach and successfully identified compounds that are already available in the market for the treatment of various other diseases, which exhibit essential properties such as anti-inflammatory drug molecules, anti-migraine compounds, anti-oxidative properties and many more (**Table 5**).

The compounds selected from virtual screening and biophysical studies have demonstrated potential against SARS-CoV-2 RdRp complex. Specifically, out of the compounds tested, seven have exhibited strong inhibitory potential against SARS-CoV-2 in *in vitro* studies. Given very promising results in *in vitro* studies, the three most promising compounds are fangchinoline, cepharanthine, and sennoside B, that can further be proceeded for follow-up validation in animal models against SARS-CoV-2. Among these, fangchinoline, a novel anti-HIV-1 drug (Wan et al., 2012), has demonstrated strong binding affinity, ranging from 7-10 kcal/mol, against catalytic and interface residues of SARS-CoV-2. It has also exhibited significant potential as a compound against the virus, with KD values of 0.67 μM and 0.18 μM for the nsp8 and nsp7 proteins, respectively in biophysical assays. Moreover, fangchinoline has shown an EC_50_ value of ∼2.80 μM, further highlighting its effectiveness against SARS-CoV-2. Interestingly, sennoside B, commonly used to treat constipation, has also shown significant antiviral action against SARS-CoV-2 with an EC_50_ value of 2.80 μM (Dreessen et al., 1981; Esposito et al., 2016; Hardcastle and Wilkins, 1970; Kon et al., 2014). Both of these compounds exhibited the highest S.I. values among all the compounds tested in *in vitro* studies against SARS-CoV-2. As a result, both compounds were selected for further investigation *in vivo* studies.

The compound cepharanthine, xifaxan, doramectin and amentoflavone, also makes H bonds and hydrophobic interactions with residues of the SARS-CoV-2 proteins via carbonyl or nitro group. These compounds have also demonstrated significant inhibitory potential against SARS-CoV-2. These molecules are known for anti-parasitic, anti-oxidative, anti-inflammatory, and anti-migraine properties as mentioned in table 5. Anti-SARS-CoV-2 activity of cepharanthine has been reported by several research groups. As per published literature, cepharanthine is speculated to act on the virus entry stage (Fan et al., 2022). In another study, Li et al., 2021 explained the potential anti-SARS-CoV-2 role of cepharanthine through reversal of the viral interference of HSF1-mediated heat shock response, hypoxia pathways, and ER stress/response to unfolded protein (Li et al., 2021). However, none of the studies stated RdRp complex or RdRp subunits as targets of cepharanthine as proposed in the current study. Moreover, the *in vivo* anti-SARS-CoV-2 activity of cepharanthine was reported by Lu et al., 2021 wherein a remarkable reduction in viral titre and lung damage was observed in cepharanthine treated mice when compared to control mice (Lu et al., 2021). Since, cepharanthine, one of the potent inhibitor of RdRp complex identified in this study has already been proven to be an effective antiviral molecule against SARS-CoV-2, these finding further validate the binding kinetics and cell based data generated in this study. Remarkably, natacyn, a compound from the natural product library, has also exhibited substantial antiviral potential against SARS-CoV-2. It is noteworthy that natacyn, which received FDA approval in 2011, is primarily used as a medication for antifungal therapy (Arora et al., 2011).

However, despite displaying favorable binding in biophysical assays, dihydroergotamine methanesulfonate, chikusetsusaponin iva, and rapamycin did not exhibit antiviral effects in *in vitro* studies possibly due to their high molecular weight (679.8 g/mol, 795.0 g/mol, and 914.2 g/mol respectively) or may be short half-life like dihydroergotamine methanesulfonate has 9 h half-life. It is worth noting that these three compounds are utilized in various medications. Dihydroergotamine methanesulfonate, for instance, is employed as an antimigraine agent (Villalon et al., 1992), while chikusetsusaponin iva is known for its antiviral, anti-inflammatory, and antithrombotic properties (Dahmer et al., 2012; Rattanathongkom et al., 2009; Xu et al., 2021). Additionally, rapamycin serves as an immunosuppressive and antitumor agent (Houchens et al., 1983; Malagelada et al., 2010).

In summary, out of the three different compound libraries, seven compounds have demonstrated significant antiviral potential against SARS-CoV-2 in the *in vitro* antiviral assays. Due to these promising results, we proceeded for *in vivo* study for the best two compounds i.e., fangchinoline and sennoside B. Thus, this study not only offers promising treatment for COVID-19 but also provides valuable structural insights for design and development of more potent analogues based on the selected compounds against SARS-CoV-2.

## Conclusion

COVID-19 is an imperative risk to global health and safety. Thus, it is crucial to halt its transmission and lower the mortality rate. Controlling the infection source and blocking the transmission pathway is crucial for removing this threat. This study describes the most promising antiviral compounds obtained from the three different libraries. Our findings from the biophysical and cell-culture based assays indicate that the fangchinoline, cepharanthine, and sennoside B compounds is postulated to have improved efficacy in tackling COVID-19. Currently, *in vivo,* studies are underway to further investigate the efficacy of these compounds. In conclusion, these compounds are considered RdRp polymerase inhibitors and pave the way for further testing against COVID-19.

## Acknowledgements

The authors are thankful to Department of Biosciences and Bioengineering, IIT Roorkee for computational facilities. RR and SC would like to thank University Grants Commission (UGC) and Council of Scientific and Industrial Research for providing financial support. SKN and AU thank Ministry of Human Resource Development (MHRD) for financial support. ST and PK acknowledges and thanks Intensification of Research in High Priority Areas (IRHPA) program of Science and Engineering Research Board (SERB), Department of Science & Technology (DST), Government of India for supporting this study (Grant No.-IPA/2020/000054). The authors are also thankful to The Director IVRI and Joint Director CADRAD IVRI for providing necessary facility to carry out work in BSL3 facility.

