## Supplementary File for "Revealing and evaluation of antivirals targeting multiple druggable sites of RdRp complex in SARS-CoV-2"

***Supplementary table 1:*** *RdRp SARS-CoV-2 protein grid box parameters for virtual screening and molecular docking*

| Proteins (Target site) | Residues | Center (Å) (X, Y, Z) | Dimensions for screening (Å) (X, Y, Z) | Dimensions for Vina (Å) (X, Y, Z) |
|---|---|---|---|---|
| **nsp12 (Catalytic Residues)** | Lys551, Arg555, Asp618, Phe620, Lys621, Cys622, Asp623, Thr680, Ser682, Gly683, Thr687, Ala688, Asn691, Ser759, Asp760, Asp761, Lys798 | 119, 117, 128 | 25, 21, 25 | 58, 66, 58 |
| **nsp12 (Interactive residues with nsp8)** | Leu270, Leu271, Tyr273, Thr324, Phe326, Pro328, Leu329,Val330, Arg331, Lys332, Val338, Pro339, Phe340, Val341, Phe368, Leu371, Tyr374, Ala375, Ala379, Met380, His381, Ala382, Ala383, Ser384, Asn386, Leu387, Leu388, Leu389, Lys391, Arg392, Val398, Ala400, Thr402, Asn403, Val405, Phe407, Pro505, Phe506, Trp509, Leu514, Tyr515, Ser518, Met666 | 116, 144, 138 | 47, 33, 34 | 126, 96, 98 |
| **nsp12 (Interactive residues with nsp7)** | Thr409, Lys411, Pro412, Gly413, Phe415, Tyr420, Glu431, Phr440, Phe442, Ala443 | 115, 102, 149 | 39, 33, 23 | 43, 94, 44 |
| **nsp8 interactive residues** | Lys79, Arg80, Val83, Thr84, Ala86, Met87, Met90, Leu91, Phe92, Met94, Leu95, Leu98, Asp99, Leu103, Asn104, Ile106, Ile107,Asn109, Ala110, Asp112, Gly113, Cys114, Val115, Pro116, Leu117, Asn118, Ile119, Ile120, Pro121, Leu122, Lys127, Leu128, Met129, Val130, Val131, Pro133, Thr141, Tyr149, Pro183 | 120, 147, 143 | 51, 49, 34 | 126, 102, 78 |
| **nsp7 interactive residues** | Lys2, Ser4, Asp5, Lys7, Cys8, Val11, Leu14, Gln18, Trp29, Val33, Asn37, Leu40, Leu41 | 124, 102, 149 | 25, 35, 23 | 36, 84, 52 |

***Supplementary table 2:*** *Binding energies (kcal/mol) of positive controls and selected molecules against the catalytic site of SARS-CoV-2 RdRp proteins*

| **S.No.** | **Ligands** | **Targeted nsp12 Catalytic Site** |
|---|---|---|
| **Positive Control** | | |
| **1.** | AMP | -6.7 |
| **2.** | CMP | -6.2 |
| **3.** | GMP | -7.0 |
| **4.** | UMP | -6.6 |
| **5.** | ATP | -6.9 |
| **6.** | CTP | -6.7 |
| **7.** | GTP | -7.5 |
| **8.** | UTP | -7.0 |
| **9.** | Remdesivir | -7.3 |
| **10.** | Sofosbuvir | -7.4 |
| **11.** | Ribavirin | -6.0 |
| **12.** | Galidesivir | -6.6 |
| **13.** | Favipiravir | -5.3 |
| **14.** | Tenofovir | -6.5 |
| **FDA Library** | | |
| **1.** | Paritaprevir | -11.4 |
| **2.** | Xifaxan | -10.5 |
| **3.** | Doramectin | -9.7 |
| **4.** | Sennoside B | -9.8 |
| **5.** | Sennoside A | -9.7 |
| **6.** | Selamectin | -9.4 |
| **7.** | Simeprevir | -9.4 |
| **8.** | Temsirolimus | -9.2 |

| | | |
|---|---|---|
| **9.** | Salvianolic acid B | -9.2 |
| **10.** | Ivermectin | -9 |
| **11.** | Avermectin B1 | -9 |
| **Natural Product Library Compounds** | | |
| **12.** | Xifaxan | -10.5 |
| **13.** | Actinomycin D | -10.3 |
| **14.** | Ligustroflavone | -10.2 |
| **15.** | O-Pentagalloylglucose | -10 |
| **16.** | Cepharanthine | -9.9 |
| **17.** | Cefoperazone acid | -9.8 |
| **18.** | Dioscin | -9.8 |
| **19.** | Punicalagin | -9.7 |
| **20.** | Echinacoside | -9.7 |
| **21.** | Rifapentine | -9.6 |
| **22.** | Amentoflavone | -9.6 |
| **23.** | Madecassoside | -9.6 |
| **24.** | Polyphyllin I | -9.6 |
| **25.** | Rosamultin | -9.5 |
| **26.** | Asiaticoside | -9.5 |
| **27.** | Tubeimoside I | -9.5 |
| **28.** | Gracillin | -9.5 |
| **29.** | Hederacoside C | -9.5 |
| **30.** | Tannic acid | -9.5 |
| **31.** | Anemoside B4 | -9.5 |
| **32.** | Forsythoside B | -9.5 |
| **33.** | Eriocitrin | -9.5 |
| **34.** | Angoroside C | -9.5 |

| | | |
|---|---|---|
| **35.** | Absinthin | -9.4 |
| **36.** | Methyl-Hesperidin | -9.4 |
| **37.** | Polyphyllin B | -9.4 |
| **38.** | Fangchinoline | -9.4 |
| **39.** | Astragaloside_IV | -9.4 |
| **40.** | Safflower Yellow | -9.4 |
| **41.** | Corilagin | -9.4 |
| **42.** | Picfeltarraenin_IA | -9.4 |
| **43.** | Oroxin_B | -9.4 |
| **44.** | Complanatuside | -9.4 |
| **45.** | Vaccarin | -9.4 |
| **46.** | Poliumoside | -9.3 |
| **47.** | Rapamycin | -9.3 |
| **48.** | Acarbose | -9.2 |
| **49.** | Chikusetsusaponin Iva | -9.2 |
| **50.** | Glycyrrhizic acid | -9.2 |
| **51.** | Hesperidin | -9.2 |
| **52.** | Kaempferol-3-O-rutinoside | -9.2 |
| **53.** | Liriope muscar baily saponins C | -9.2 |
| **54.** | Naringin | -9.2 |
| **55.** | Rifampin | -9.2 |
| **56.** | Ginsenoside Rg1 | -9.2 |
| **57.** | Dipsacoside B | -9.2 |
| **58.** | Rubusoside | -9.2 |
| **59.** | Phytolaccagenin | -9.2 |
| **60.** | Veratrine | -9.2 |

| | | |
|---|---|---|
| **61.** | Eptifibatide Acetate | -9.1 |
| **62.** | Salvianolic acid B | -9.1 |
| **63.** | Pectolinarin | -9.1 |
| **64.** | Narirutin | -9.1 |
| **65.** | Alisol A | -9.1 |
| **66.** | Aminophylline | -9 |
| **67.** | Digoxin | -9 |
| **68.** | Diosmin | -9 |
| **69.** | Linarin | -9 |
| **70.** | Lypressin Acetate | -9 |
| **71.** | Natacyn | -9 |
| **72.** | Polyphyllin VII | -9 |
| **73.** | Fangchinoline | -9 |
| **74.** | Avermectin B1 | -9 |
| **75.** | Maduramycin Ammonium | -9 |
| **76.** | Verbascoside | -9 |
| **77.** | Kaempferitrin | -9 |
| **78.** | Epimedin A | -9 |
| **79.** | Narcissoside | -9 |
| **80.** | Pedunculoside | -9 |
| **81.** | Sorafenib | -9 |
| **82.** | Taxifolin 7-rhamnoside | -9 |
| **LOPAC** | | |
| **83.** | Dihydroergotamin methanesulfonate | -10 |
| **84.** | WIN 62577 | -9.7 |
| **85.** | Eptifibatide acetate | -9.5 |
| **86.** | KT185 | -9.3 |

| | | |
|---|---|---|
| **87.** | Dihydroouabain | -9.1 |
| **88.** | KT203 | -9.1 |
| **89.** | Stevioside | -9.1 |
| **90.** | Ivermectin | -9 |

***Supplementary table 3:*** *Binding energies (kcal/mol) of selected molecules from virtual screening against the nsp12 interface residues of SARS-CoV-2 proteins*

| **S.No.** | **Ligands** | **nsp12 interface residues against** | | **Targeted interface residues** | |
|---|---|---|---|---|---|
| | | **nsp8** | **nsp7** | **nsp8** | **nsp7** |
| **FDA Library from Selleckchem** | | | | | |
| **1.** | Sennoside B | -10.4 | -8.7 | -8.0 | -7.2 |
| **2.** | Sennoside A | -10.4 | -8.7 | -7.8 | -7.8 |
| **3.** | Paritaprevir | -10.3 | -9.6 | -8.2 | -7.6 |
| **4.** | Doramectin | -10.1 | -8.5 | -7.9 | -7.2 |
| **5.** | Selamectin | -10.1 | -8.3 | -7.5 | -7.0 |
| **6.** | Simeprevir | -9.6 | -8.8 | -7.3 | -7.0 |
| **7.** | Xifaxan | -9 | -9 | -7.4 | -7.2 |
| **Natural Product Library Compounds from Selleckchem** | | | | | |
| **8.** | Amentoflavone | -10.6 | -9.2 | -7.8 | -7.6 |
| **9.** | Phytolaccagenin | -9.6 | -8.7 | -6.8 | -6.1 |
| **10.** | Natacyn | -9.5 | -8.2 | -7.1 | -7.0 |
| **11.** | Oroxin B | -9.5 | -8.2 | -7.1 | -6.0 |
| **12.** | Rapamycin | -9.5 | -8.1 | -7.8 | -7.0 |
| **13.** | Chikusetsusaponin Iva | -9.5 | -8.0 | -7.8 | -7.4 |
| **14.** | Veratrine | -9.3 | -8.1 | -9.0 | -7.0 |
| **15.** | Cepharanthine | -9.2 | -9.3 | -8.0 | -7.2 |
| **16.** | Alisol A | -9.2 | -8.4 | -6.7 | -6.0 |
| **17.** | Absinthin | -9 | -9.1 | -8.2 | -7.0 |

| | | | | | |
|---|---|---|---|---|---|
| **18.** | Fangchinoline | -9 | -8.9 | -7.5 | -7.0 |
| **19.** | Astragaloside IV | -9 | -8.2 | -7.8 | -6.3 |
| **LOPAC from Sigma** | | | | | |
| **20.** | Dihydroergotamine | -10.4 | -8.8 | -8.9 | -7.2 |
| **21.** | KT203 | -9.7 | -8.3 | -7.6 | -7.2 |
| **22.** | KT185 | -9.6 | -9.7 | -9.0 | -7.3 |
| **23.** | WIN62577 | -9.3 | -8.3 | -7.3 | -7.0 |

### Supplementary Figures:

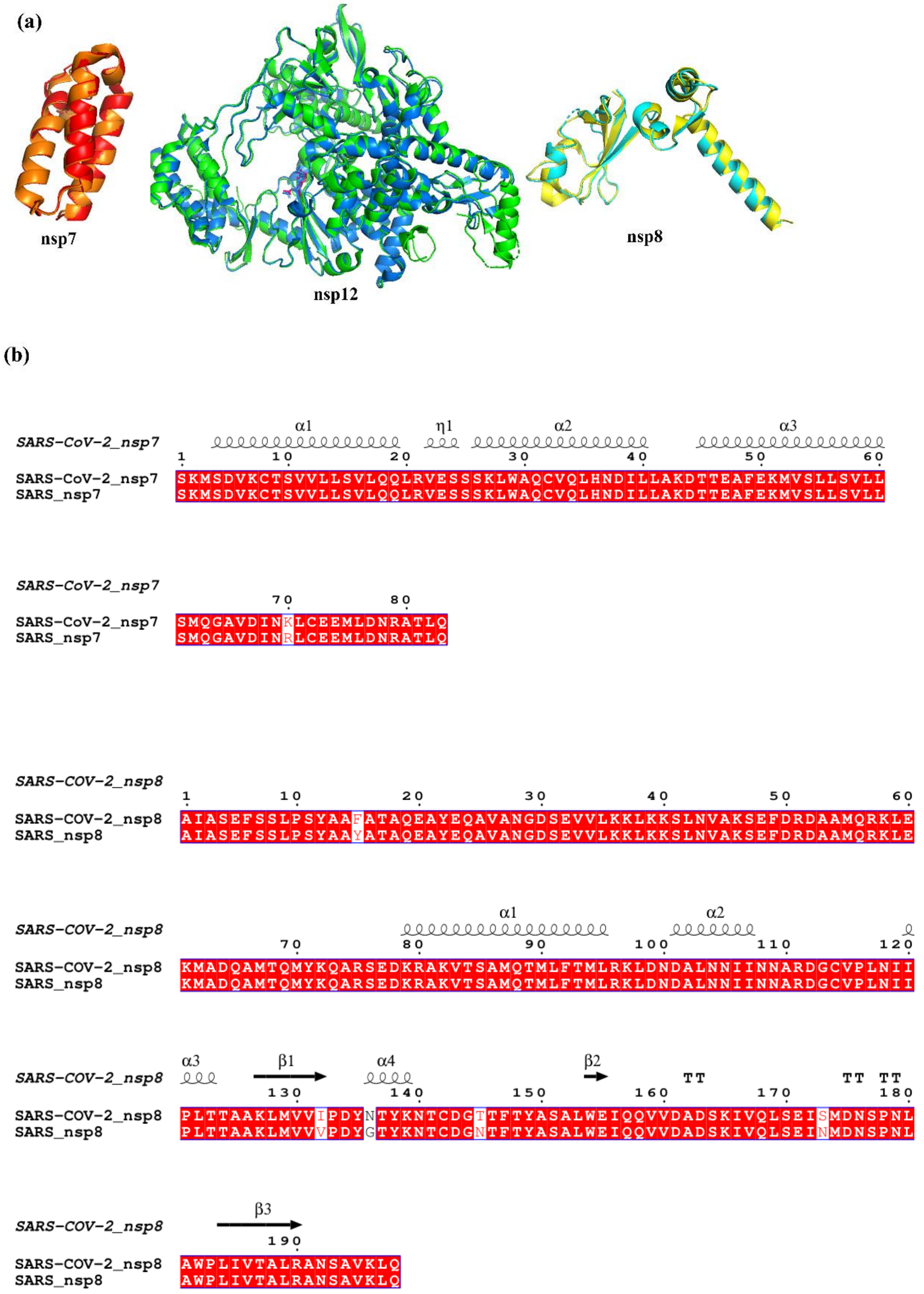

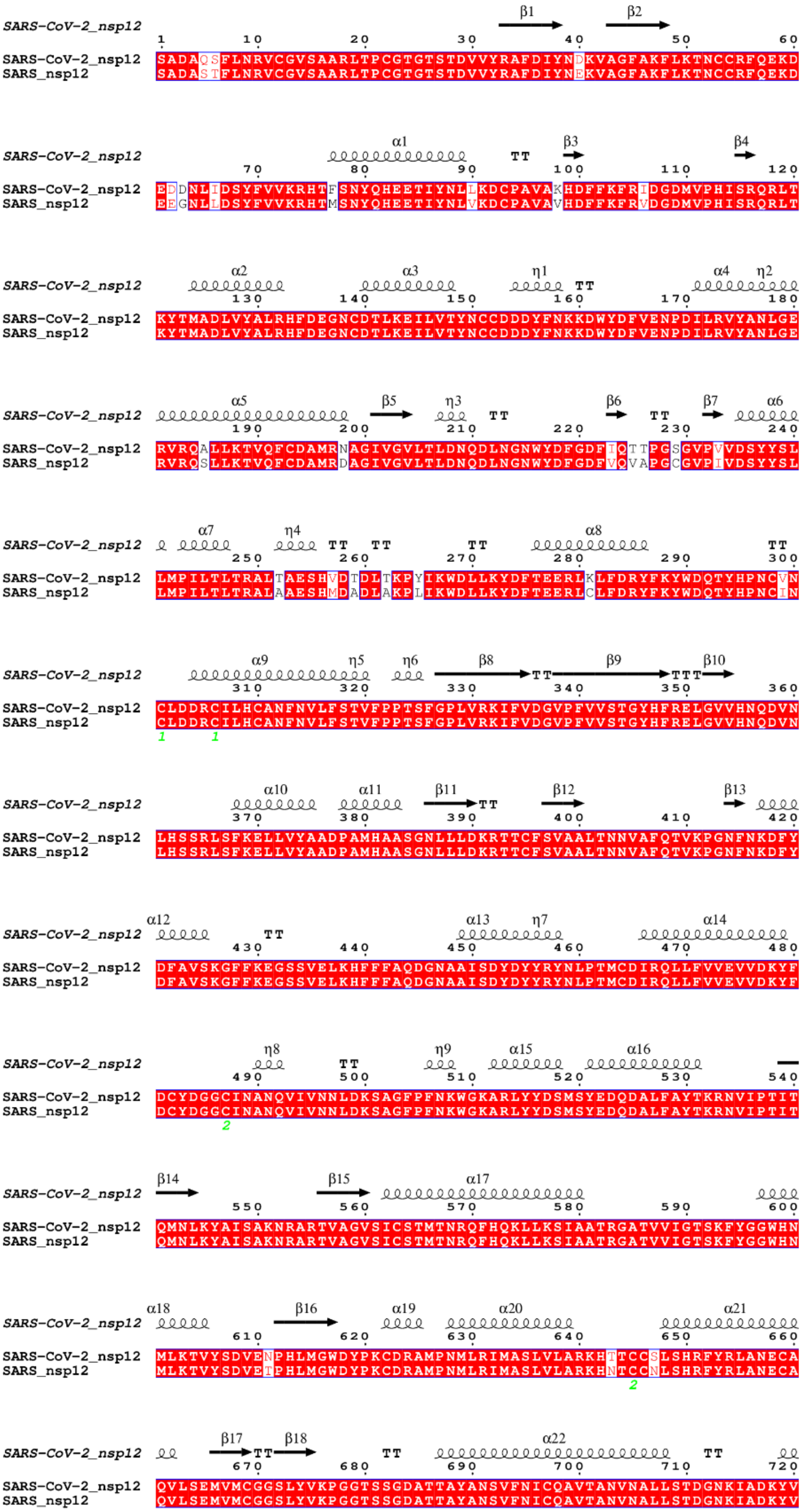
SARS-CoV-2_nsp12
β1 β2
SARS-CoV-2_nsp12 SADAQSFLNRVCGVSAARLTPCGTGTSTDVVYRAFDIYNDKVAGFAKFLKTNCCRFQEKD
SARS_nsp12 SADASTFLNRVCGVSAARLTPCGTGTSTDVVYRAFDIYNEKVAGFAKFLKTNCCRFQEKD
α1 TT β3 β4
SARS-CoV-2_nsp12 EDDNLIDSYFVVKRHTFSNYQHEETIYNLLKDCPAVAKHDFFKFRIDGDMVPHISRQRLT
SARS_nsp12 EEGNLLDSYFVVKRHTMSNYQHEETIYNLVKDCPAVAVHDFFKFRVDGDMVPHISRQRLT
α2 α3 η1 TT α4 η2
SARS-CoV-2_nsp12 KYTMADLVYALRHFDEGNCDTLKEILVTYNCCDDDYFNKKDWYDFVENPDILRVYANLGE
SARS_nsp12 KYTMADLVYALRHFDEGNCDTLKEILVTYNCCDDDYFNKKDWYDFVENPDILRVYANLGE
α5 β5 η3 TT β6 TT β7 α6
SARS-CoV-2_nsp12 RVRQALLKTVQFCDAMRNAGIVGVLTLDNQDLNGNWYDFGDFIQTTPGSGVPVVDSYYSL
SARS_nsp12 RVRQSLLKTVQFCDAMRDAGIVGVLTLDNQDLNGNWYDFGDFVQVAPGCGVPIVDSYYSL
α7 η4 TT TT TT α8 TT
SARS-CoV-2_nsp12 LMPILTLTRALTAESHVDTDLTKPYIKWDLLKYDFTEERLKLFDRYFKYWDQTYHPNCVN
SARS_nsp12 LMPILTLTRALAAESHMDADLAKPLIKWDLLKYDFTEERLCLFDRYFKYWDQTYHPNCIN
α9 η5 η6 β8 TT β9 TTT β10
SARS-CoV-2_nsp12 CLDDRCILHCANFNVLFSTVFPPTSFGPLVRKIFVDGVPFVVSTGYHFRELGVVHNQDVN
SARS_nsp12 CLDDRCILHCANFNVLFSTVFPPTSFGPLVRKIFVDGVPFVVSTGYHFRELGVVHNQDVN
1 1
α10 α11 β11 TT β12 β13
SARS-CoV-2_nsp12 LHSSRLSFKELLVYAADPAMHAASGNLLLDKRTTCFSVAALTNNVAFQTVKPGNFNKDFY
SARS_nsp12 LHSSRLSFKELLVYAADPAMHAASGNLLLDKRTTCFSVAALTNNVAFQTVKPGNFNKDFY
α12 TT α13 η7 α14
SARS-CoV-2_nsp12 DFAVSKGFFKEGSSVELKHFFFAQDGNAAISDYDYYRYNLPTMCDIRQLLFVVEVVDKYF
SARS_nsp12 DFAVSKGFFKEGSSVELKHFFFAQDGNAAISDYDYYRYNLPTMCDIRQLLFVVEVVDKYF
η8 TT η9 α15 α16
SARS-CoV-2_nsp12 DCYDGGCINANQVIVNNLDKSAGFPFNKWGKARLYYDSMSYEDQDALFAYTKRNVIPTIT
SARS_nsp12 DCYDGGCINANQVIVNNLDKSAGFPFNKWGKARLYYDSMSYEDQDALFAYTKRNVIPTIT
2
β14 β15 α17
SARS-CoV-2_nsp12 QMNLKYAISAKNRARTVAGVSICSTMTNRQFHQKLLKSIAATRGATVVIGTSKFYGGWHN
SARS_nsp12 QMNLKYAISAKNRARTVAGVSICSTMTNRQFHQKLLKSIAATRGATVVIGTSKFYGGWHN
α18 β16 α19 α20 α21
SARS-CoV-2_nsp12 MLKTVYSDVENPHLMGWDYPKCDRAMPNMLRIMASLVLARKHTTCCSLSHRFYRLANECA
SARS_nsp12 MLKTVYSDVETPHLMGWDYPKCDRAMPNMLRIMASLVLARKHNTCCNLSHRFYRLANECA
2
β17 TT β18 TT α22 TT
SARS-CoV-2_nsp12 QVLSEMVMCGGSLYVKPGGTSSGDATTAYANSVFNICQAVTANVNALLSTDGNKIADKYV
SARS_nsp12 QVLSEMVMCGGSLYVKPGGTSSGDATTAYANSVFNICQAVTANVNALLSTDGNKIADKYV

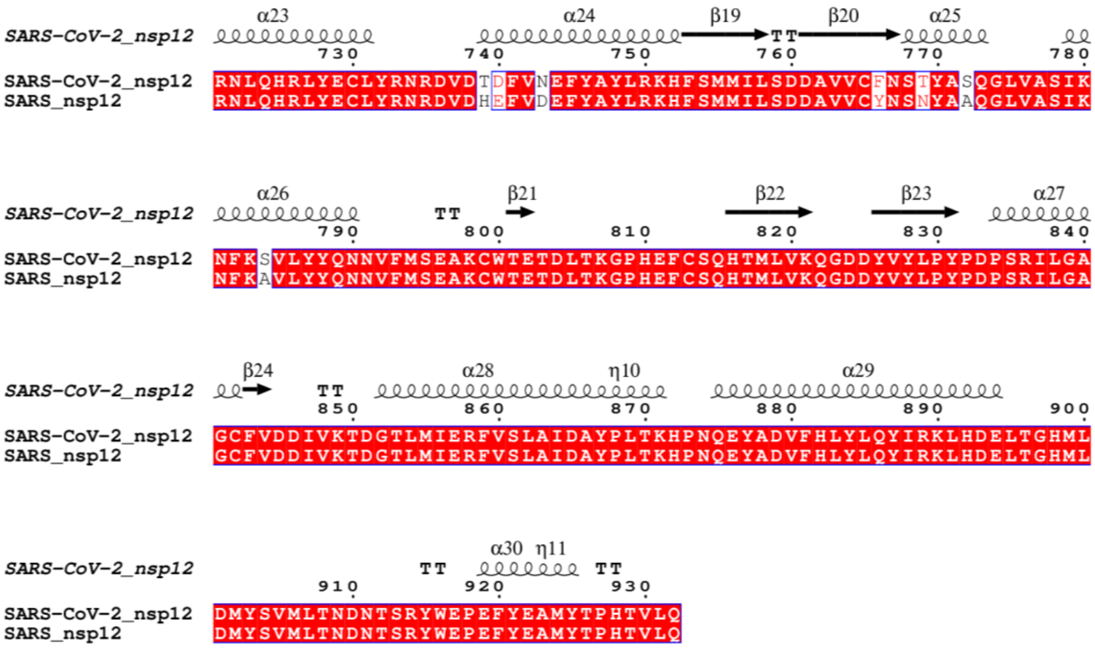


***Supplementary figure 1**: (a) Superimposed protein structures of SARS-CoV-2 and SARS RdRp complex proteins (nsp7, nsp12 and nsp8) using PyMOL (PDB ID:6M71). Red-orange for nsp7, green-blue for nsp12, cyan-blue of nsp8 proteins of SARS-COV-2 and SARS respectively. RMSD values for nsp7, nsp12 and nsp8 are 0.399, 0.482, and 0.639 respectively. (b) Multiple sequence alignment of nsp7, nsp8 and nsp12 of SARS and SARS-CoV-2 by Cluster Omega and visualize by ESPript3. Red color indicates conserved residues.*

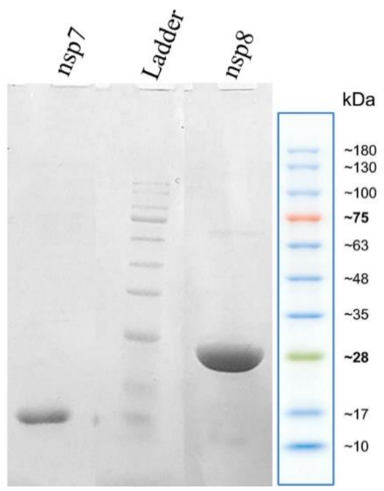


***Supplementary figure 2:*** *Purification of nsp7 and nsp8 by Ni$^{+2}$-NTA affinity chromatography. SDS-PAGE profile of purified nsp8 (24 kDa), nsp7 (10.4 kDa). The protein ladder is from 10 to 180 kDa.*

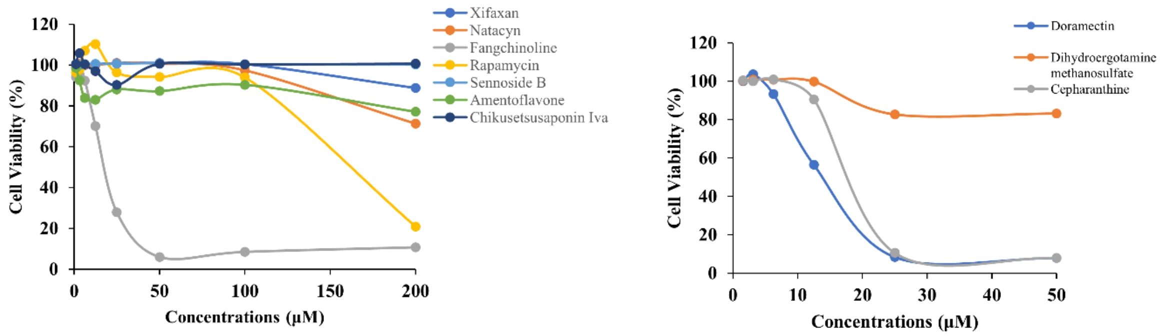


***Supplementary figure 3:*** *Cytotoxicity of selected compounds in Vero cells. Vero cells were exposed to a series of two-fold diluted concentrations of the compounds ranging from 200 μM -1.56 μM and incubated at 37 °C in 5% $CO_2$ for 48 hours for MTT assay (depicted in cell color). The relative activity of Vero cells treated with 0.1% DMSO was considered as cell control or 100% viability, and the cytotoxicity is expressed as the percentage of cell activity relative to the DMSO-treated cells. Each data point represents three independent replicate experiments.*

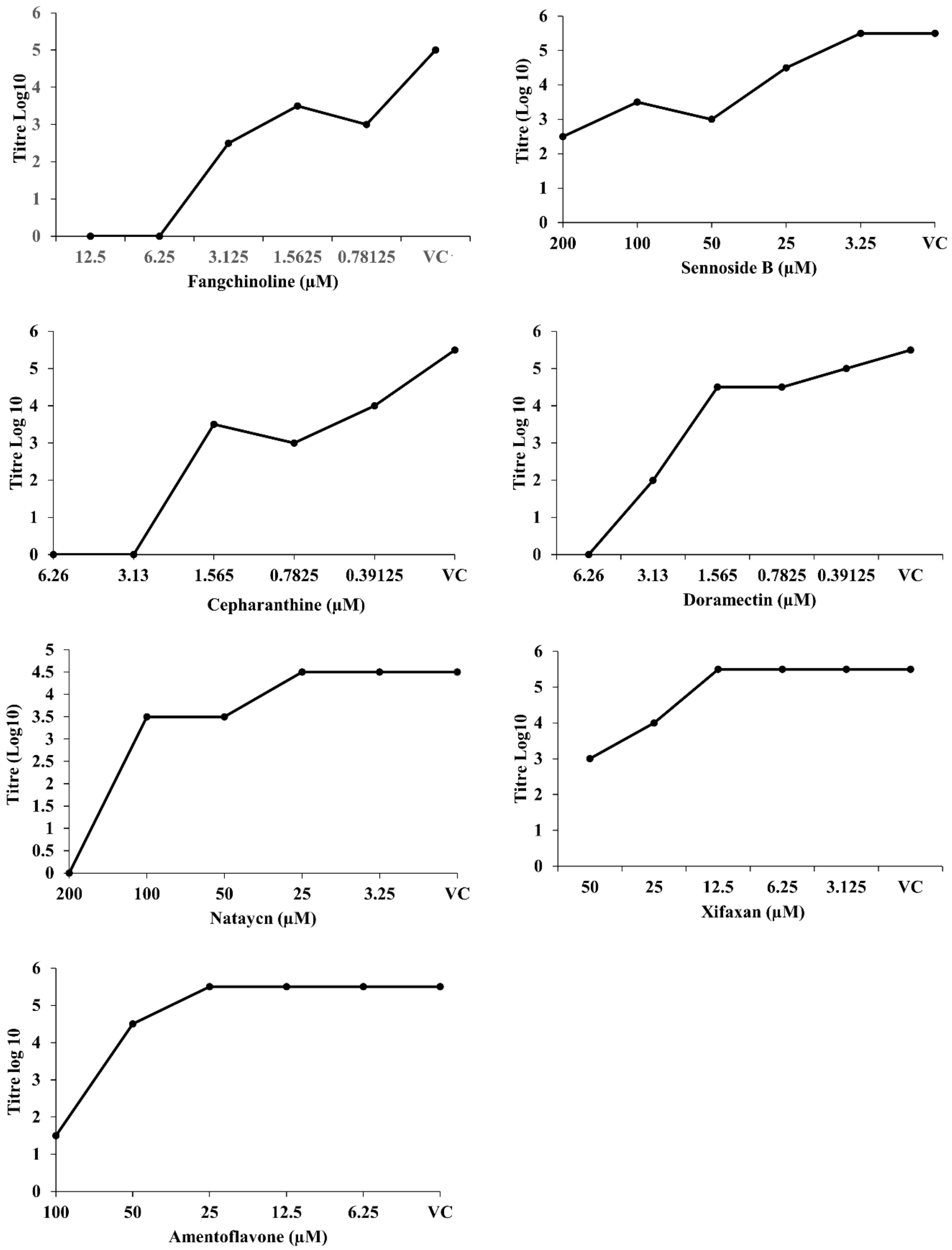


***Supplementary Figure 4**: $TCID_{50}$ growth curves of selected compounds against SARS-CoV-2. Vero cells were seeded into a 96-well plate and incubated at 37 °C with 5 % $CO_2$ until reaching confluency. The freeze-thawed cell lysate of infected and compounds treated cells were collected at 48 hpi of the antiviral assessment was subjected to ten-fold serial dilutions for 3-4 days at 37 °C and 5 % $CO_2$, with daily monitoring for the appearance of a CPE. Mock treated*

*and infected Vero cell were used as a vehicle control (VC) and samples containing only culture media were used as a cell control (CC).*
